# The structural role of bacterial eDNA in the formation of biofilm streamers

**DOI:** 10.1101/2021.07.26.453754

**Authors:** Eleonora Secchi, Giovanni Savorana, Alessandra Vitale, Leo Eberl, Roman Stocker, Roberto Rusconi

## Abstract

Across diverse habitats, bacteria are mainly found as biofilms, surface-attached communities embedded in a self-secreted matrix of extracellular polymeric substances (EPS), which enhances bacterial resistance to antimicrobial treatment and mechanical stresses. In the presence of flow and geometric constraints such as corners or constrictions, biofilms take the form of long, suspended threads known as streamers, which bear important consequences in industrial and clinical settings by causing clogging and fouling. The formation of streamers is thought to be driven by the viscoelastic nature of the biofilm matrix. Yet, little is known about the structural composition of streamers and how it affects their mechanical properties. Here, using a microfluidic platform that allows growing and precisely examining biofilm streamers, we show that extracellular DNA (eDNA) constitutes the backbone and is essential for the mechanical stability of *Pseudomonas aeruginosa*’ s streamers. This finding is supported by the observations that DNA-degrading enzymes prevent the formation of streamers and clear already formed ones, and that the antibiotic ciprofloxacin promotes their formation by increasing the release of eDNA. Furthermore, using mutants for production of the exopolysaccharide Pel, an important component of *P. aeruginosa*’ s EPS, we reveal a new, although indirect role of Pel, in tuning the mechanical properties of the streamers. Taken together, these results highlight the importance of eDNA and of its interplay with Pel in determining the mechanical properties of *P. aeruginosa* streamers, and suggest that targeting the composition of streamers can be an effective approach to control the formation of these biofilm structures.

## Introduction

The extracellular matrix confers stability to biofilms (1, 2) and protects the bacterial community against chemical and mechanical insults (3, 4). This protected environment makes biofilm bacteria a major cause of chronic infections in clinical environments and of clogging in industrial flow systems (5–7). The diverse biopolymers composing the extracellular matrix form the three-dimensional scaffold of the biofilm and are responsible for both adhesion to the surface and cohesion within the biofilm. The EPS include polysaccharides, proteins, extracellular DNA (eDNA) and lipids; however, the composition can vary greatly depending on the microorganisms present, the shear forces experienced, temperature and nutrient availability (3). The chemical composition and the resulting intermolecular interactions between EPS components drive the self-assembly of the components and determine the structure and the mechanical properties of the biofilm network (8–11). The rheological signature of biofilms is a viscoelastic response to external forces, characterized by an instantaneous elastic deformation followed by a viscous flow, which allows dissipation of long-lasting stresses without structural failure (12). This rheological behavior is key to biofilm persistence in flow (13, 14) and allows biofilms to adapt their architecture to colonize very diverse environments (15–17).

Biofilm streamers are perhaps the most profound example of biofilm structural specialization in response to environmental conditions (18–20). Streamers take the form of long filamentous structures suspended within a flow, which stem from the interaction with the physical environment and increase the carrying capacity of biofilms in natural ecosystems (21). Indeed, their localization within the bulk fluid flow, in contrast to the thin-film morphology of surface-associated biofilms, confers streamers a greater impact on the uptake of microorganisms and debris flowing by and clogging (22, 23). Structures morphologically similar to streamers can be formed also by abiotic materials in flow, as it has been shown by flowing a suspension of polyacrylamide and polystyrene particles through a porous structure (24). Aggregation processes and the viscous nature of the polymeric matrix play a crucial role in the formation of these filamentous structures (25, 26). These observations suggest that streamers formation is driven by multiple factors, including the local three-dimensional hydrodynamic profile and EPS mechanical properties, in addition to bacterial physiology. Yet, to date, little is known about the impact of EPS matrix composition on streamers morphology and rheology.

The opportunistic pathogen *Pseudomonas aeruginosa* is one of the best-characterized model organisms in the study of biofilm development (27–31). It forms biofilms in diverse habitats, including soil, rivers, medical devices and human organs (6, 32, 33), with different architectures: in the surface-attached morphology, *P. aeruginosa* biofilms often show a characteristic mushroom-like shape (34, 35), whereas in a flowing fluid they can form streamers (18, 19, 22). *P. aeruginosa* can synthesize three exopolysaccharides: alginate, Pel and Psl (36). Alginate production is typical of mucoidal strains isolated from patients with chronic pulmonary infections and is usually lost upon domestication (27). Most environmental and clinical isolates produce the polysaccharide Pel and some also produce the polysaccharide Psl (16, 36–39). Psl promotes early colony formation, increases elasticity, and strengthens the scaffold of biofilms, whereas Pel promotes adhesion among cells and to surfaces, decreases biofilm viscosity thereby increasing the ability to spread on surfaces (16), and provides protection against antibiotics (38). In strains deficient in the Psl gene cluster, such as PA14, Pel is the primary exopolysaccharide and forms the scaffold of the entire biofilm (28, 29, 38, 40).

eDNA is a functional component of *P. aeruginosa* biofilms (41–43). eDNA was originally considered a minor component, mostly relevant as a gene pool to be exploited for horizontal gene transfer (44), but its contribution to biofilm formation has been recently re-evaluated (42, 43, 45). The presence of eDNA has also been documented in single- and multi-species biofilms formed in different environments (34, 46–55) by *Staphylococcus aureus* (46, 48, 49, 56), *Myxococcus xanthus* (50), *Burkholderia cenocepacia* (51), *Staphylococcus epidermidis* (48, 52), *Bacillus subtilis* (53) and in mixed environmental samples (47, 54, 55). eDNA is released by cell death and lysis, which mainly occur in the interior of biofilms (e.g., the stalk of the mushrooms) (41, 57, 58), and can be induced by exogenous stresses, such as antibiotic treatment (57). Thanks to the binding capability of the highly charged DNA molecule, eDNA protects the community from antimicrobials and antibiotics (59, 60). Additionally, eDNA promotes cell–cell adhesion (41) and biofilm self-organization (61), and enables the formation of a stable biofilm structure, as indirectly demonstrated by the degradation of biofilms by treatment with DNase I (42, 62), a non-specific nuclease that cleaves DNA. Yet, a mechanistic description of the impact of eDNA on biofilm architecture is missing.

Lectin staining has shown that Pel, which is composed of positively charged amino sugars, binds eDNA in the stalk of the mushroom structures in mature biofilms, conferring stability. The ionic-binding mechanism between Pel and eDNA is pH-dependent, since Pel is cationic until the isoelectric point at pH = 7.3, where Pel carries no charge (34). This suggests that chemical heterogeneity within a biofilm (63) can influence Pel’s ability to bind to eDNA and consequently its localization within the biofilm structure. To date, eDNA-Pel co-localization and cross-linkage have been investigated in the mushroom architecture of surface-attached biofilms (34), but the structural and rheological implications of the eDNA-Pel interaction have not been studied.

In this work, we analyze the distinct role of eDNA and Pel in the formation of *P. aeruginosa* streamers and show how their interaction tunes their rheological properties. We show that eDNA is the constitutive element of the backbone of streamers, essential for their formation and stability. This finding allows controlling the formation of streamers. Treatment with DNase I, a non-specific nuclease that cleaves DNA, completely suppresses their formation and clears already formed streamers in the wild-type strain. The efficacy of the treatment is reduced in Pel-overproducing strains, due to the protecting action of Pel. On the other hand, sub-lethal concentrations of ciprofloxacin, an antibiotic known to induce cell lysis (57), can paradoxically stimulate the formation of streamers. This effect is due to the release of bacterial eDNA in the flowing solution that is captured by the streamers. By varying the composition of the EPS using Pel-mutant strains, we show that an increase in the Pel content determines an increase in the elastic modulus and the viscosity of the matrix, indicating that eDNA-Pel interaction results in stiffer streamers. Taken together, these findings provide a mechanistic understanding of the structural role of different matrix components in the formation of *P. aeruginosa* biofilm streamers.

## Results

### eDNA represents the backbone of biofilm streamers

Obstacles in a flow field act as a tethering point for the formation of biofilm streamers. This scenario, which occurs for example in porous media, in the human body and filters, can be recapitulated in its simplest form by isolated pillars in a microfluidic channel. In our microfluidic platform, reproducible formation of streamers could be achieved on 50-μm-diameter, isolated pillars (Fig. *1A*; Methods) exposed to a constant flow of a diluted suspension of *P. aeruginosa* PA14. Fluorescence and phase-contrast video microscopy were used to characterize the dynamics and composition of streamers during their formation (Fig. 1 *B-D*). The first strands formed after 3–5 h, and after about 10–15 h they had become millimeter-long stable filaments (Fig. 1*B*). Around each pillar, we observed the formation of two symmetric streamers, tethered on the side of the pillar and suspended at channel mid-depth (Fig. 1*A*). At the flow velocities used in our experiments, bacterial cells, in addition to colonizing the inner walls of the microfluidic channel, attached to the windward side of the pillar (64) and EPS accumulated around the curved pillar surface (18) before being extruded by downstream flow (26). Biomass accumulation is recognized as the dominant mechanism for streamer initiation (21) and is promoted by the secondary flow around the curved surfaces (18), while the extrusion process is favored by the extensional component of the flow downstream of the pillar.

**Figure 1.**
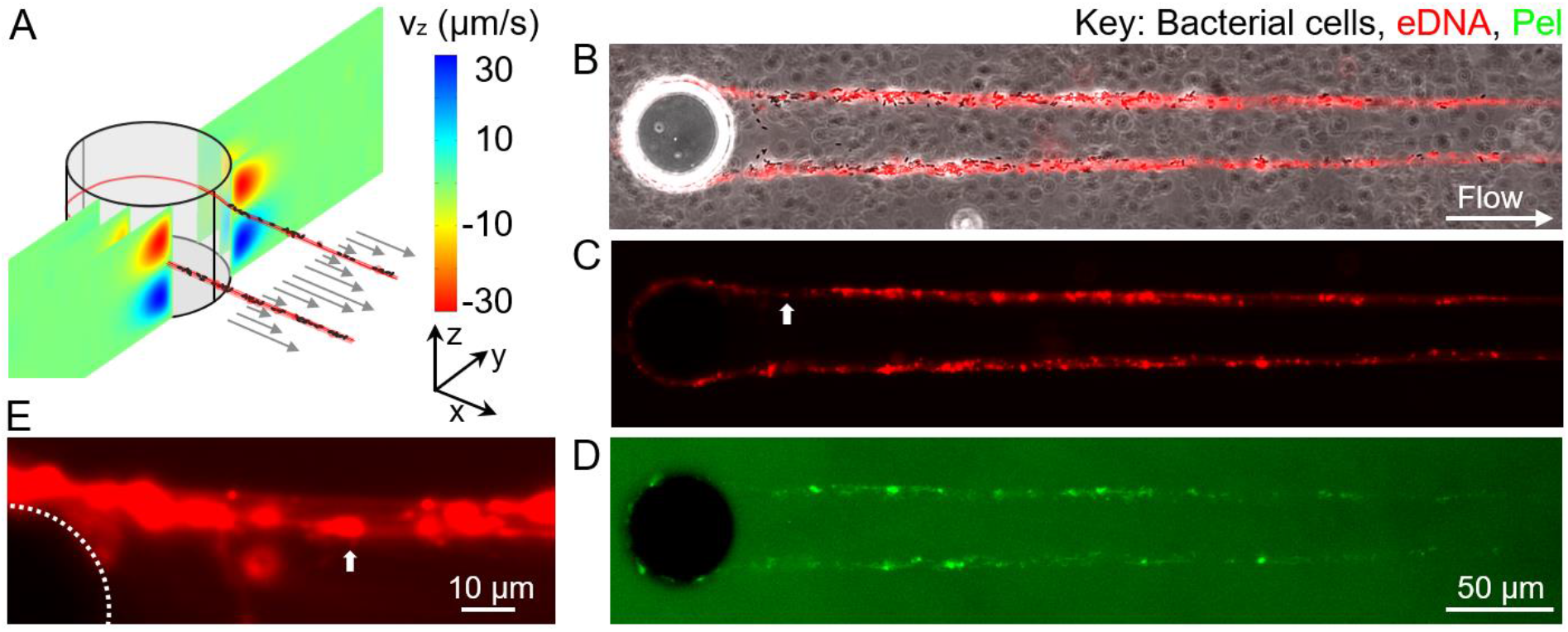
eDNA is a major component of PA14 biofilm streamers. (*A*) Schematic of a biofilm streamer formed around a pillar, due to the combined action of the main flow (gray arrows) and the secondary flow (color scale). The *x-* (gray arrows) and the *z*-component (color scale) of the flow velocity around a 50-μm pillar were numerically computed (COMSOL Multiphysics) for a mean flow velocity of 500 μm/s. (*B–D*) Phase-contrast and fluorescence images of a portion of the biofilm streamers formed by *P. aeruginosa* PA14 WT cells attached to a 50-μm pillar within a microfluidic channel after 20 h of continuous flow of a dilute bacterial suspension at *U* = 2 mm/s. Images were taken at channel mid-depth. Bacterial cells were imaged in phase contrast (*B*), eDNA was stained using red-fluorescent PI (2 μg/mL; *B*, *C, E*) and Pel was stained using green-fluorescent WFL (50 μg/mL; *D*).

This experimental system allowed us to identify the spatial localization of different components of streamers: bacterial cells were visualized with phase-contrast microscopy (Fig. 1*B*), while eDNA and Pel were visualized by epifluorescence microscopy after staining with propidium iodide (PI) (Fig. 1*C*) and *Wisteria floribunda* lectin (WFL) (Fig. 1*D*), respectively (Methods). Cell aggregates were mainly located in the first 200 μm downstream from the pillar, whereas further downstream we mainly observed single bacteria embedded in the streamers (Fig. 1*B*). Both Pel and eDNA were present on the filament: eDNA was homogeneously distributed along the streamer (Fig. 1*B*, *C*), whereas Pel was mainly co-localized with cell aggregates, resulting in a higher abundance in the region of the filament closer to the pillar (Fig. 1*D*). We observed a continuous red PI-stained thread in all the streamers, suggesting that bundles of eDNA connect the bacterial cells within the streamers (Fig. 1*C, E*). The continuous distribution of eDNA within streamer filaments indicates that eDNA formed the backbone structure of the streamers (Fig. 1*C*), whereas Pel was localized in micrometer-sized patches, mainly located in the cells’ clusters (Fig. S1, Fig. S2).

### The morphology and mechanical properties of streamers depend on Pel abundance

Pel is not essential for the formation of streamers, but it has an impact on their morphology. The role of Pel was determined by comparing the streamers formed by the wild-type strain (PA14 WT), a mutant strain lacking the ability to produce Pel (PA14 *ΔpelE*), and a Pel overproducer strain (PA14 *ΔwspF*). All three strains produced millimeter-long streamers, on the same timescale (Fig. 2 *A–C*). WFL staining confirmed that PA14 *ΔpelE* streamers lacked Pel (Fig. 2*F*, Fig. S1), while Pel-overproducer streamers showed a tenfold increase in the quantity of Pel compared to WT (Fig. 2*F*, Fig. S2). Pel-deficient streamers were on average 3.03 mm ± 0.03 mm long after 22 h, marginally (~10%) longer than WT streamers and approximately twice as long as Pel-overproducer streamers, which reached an average length of 1.47 mm ± 0.28 mm (Fig. 2*D*). Differences in Pel content also affected the average diameter of streamers, which we measured at different distances from the pillar (Material and Methods). After 22 h, WT streamers display a diameter *d*_150_ = 10 μm ± 0.2 μm in the vicinity of the pillar and *d*400 = 5 μm ± 0.3 μm in the downstream region. Compared to WT streamers, after 22 h, Pel-deficient streamers showed a diameter *d*_150_ = 8.4 μm ± 0.2 μm (~15%) in the vicinity of the pillar, which is 15% smaller compared to WT (Fig. 2*E*). Instead, the diameter of Pel-overproducer streamers near the pillar was comparable with the WT one after 15 h (*d*_150_ = 8.9 μm ± 0.3 μm for WT and *d*_150_ = 8.3 μm ± 1.2 μm for *ΔwspF*) and double that of WT streamers after 22 h (*d*_150_ = 10 μm ± 0.2 for WT and *d*_150_ = 18.6 μm ± 1.6 μm for *ΔwspF*). The trend is inverted in the downstream portion of the filament, where the diameter of the Pel-overproducer strain was approximately half that of WT streamers (Fig. 2*E*). Data on streamers’ geometry are quantified over 25 replicates. Cell aggregates were also mainly located in the vicinity of the pillar (Fig. 2*A*–*C*). Pel-deficient streamers (Fig. 2*A*) had substantially fewer and smaller cell clusters than WT streamers (Fig. 2*B*), while in Pel-overproducer streamers bacterial aggregates were larger and more abundant (Fig. 2*C*).

**Figure. 2.**
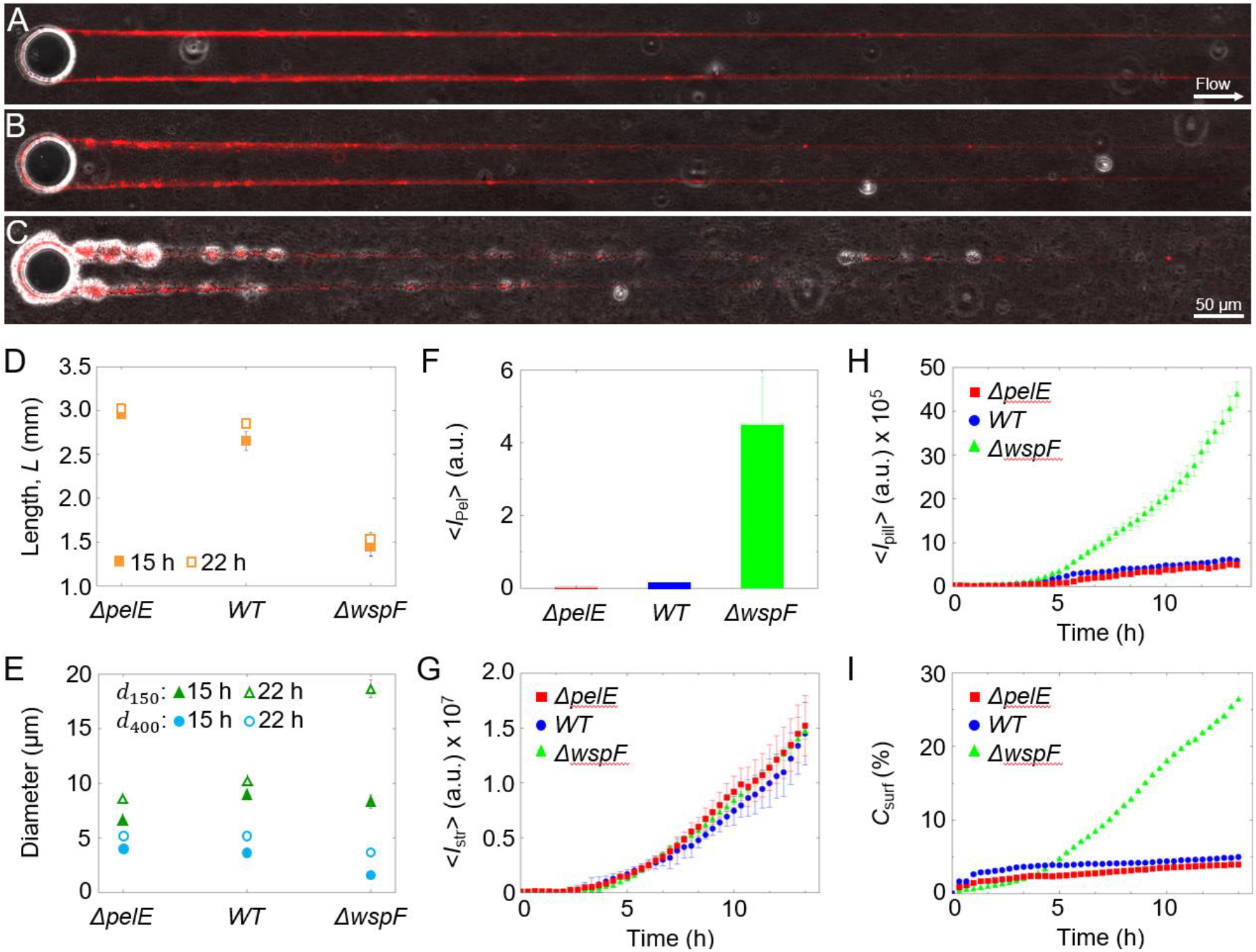
Pel affects the morphology of PA14 biofilm streamers. (*A*-*C*) Overlaid phase-contrast (bacterial cells) and red fluorescence (eDNA) images of the biofilm streamers formed by *P. aeruginosa* PA14 Pel-deficient (*ΔpelE*) (*A*), wild-type (WT) (*B*) and Pel-overproducing (*ΔwspF*) (*C*) cells attached to a 50-μm pillar after 14 h of continuous flow of a dilute bacterial suspension at *U* = 2 mm/s. (*D*) Length of streamers after 15 h (filled symbols) and 22 h (open symbols) of continuous flow, measured from fluorescence images for *ΔpelE*, WT, and *ΔwspF* strains. (*E*) Diameter of streamers in the vicinity of the pillar (150 μm from the pillar, *d*_150_) (green triangles) and in the downstream region (400 μm from the pillar, *d*_400_) (blue circles) after 15 h (filled symbols) and 22 h (open symbols) of continuous flow, measured from fluorescence images for the three strains. (*F*) Green fluorescence intensity of streamers, 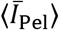, corresponding to the eDNA content, for the three strains after 24 h. (*G*) Red fluorescence intensity of streamers, 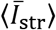, corresponding to the eDNA content, as a function of time (red squares, *ΔpelE*; blue circles, WT; green triangles, *ΔwspF*). (*H*) Red fluorescence intensity measured on the pillar surface, 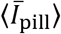, as a function of time for the same suspensions as *F*. (*I*) Coverage of the horizontal surface of the channel, *C*_surf_, as a function of time for the same suspensions as *F* and *H*. Error bars represent standard error of the mean.

Despite the differences in morphology associated with the different amounts of Pel, the quantity of eDNA per unit length of the streamer, 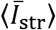, as assessed by the red fluorescence intensity of PI staining (Material and Methods, Fig. S3), was comparable in the three strains and showed the same, increasing trend with time (Fig. 2*G*). This trend is linear from 5 h onward (Fig. 2*G*) and is consistent with the previously proposed filtration model (21, 22, 26), which attributes the streamer’s growth to the addition of mass captured by the streamer from the flowing fluid. This dynamics contrasts with the differences in biofilm growth on the surface among the three strains, which was determined by analyzing the colonization of the horizontal surface of the microfluidic channel and the vertical surface of the pillar. In terms of surface coverage, quantified as the fraction *C*_surf_ of the surface covered by bacteria, the Pel-deficient strain shows the lowest surface growth, with *C*_surf_ < 4% after 14 h (Fig. 2*I*), closely followed by the WT (*C*_surf_ = 5%). In line with this observation, WT and *ΔpelE* showed limited colonization of the pillar surface compared to the *ΔwspF* mutant, as quantified by the red fluorescence intensity on the pillar, 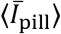 (Fig. 2*H*). The Pel-overproducing strain displayed greater surface growth, leading to five times higher surface coverage (Fig. 2*I*) and seven times higher biomass accumulation on the pillar (Fig. 2*H*) in comparison with the WT strain.

Pel not only affects the morphology of the streamers, but also their viscoelastic behavior. We studied streamer rheology *in situ* using a newly developed technique that relies on measuring the deformation of the filaments in response to a controlled variation of fluid shear (manuscript in preparation) (Methods). Specifically, streamers grown at a flow velocity of 2 mm/s were deformed by suddenly exposing them to a flow velocity of 4 mm/s for 5 min. We observed that streamers underwent a deformation typical of viscoelastic materials: on a timescale of seconds, they displayed an elastic deformation, which was recovered once flow velocity was lowered, followed by a slow viscous deformation on a timescale of minutes (Fig. S4). All three strains exhibited this viscoelastic deformation, however, the elastic modulus *E* and the effective viscosity *η* associated with the deformation depended on the Pel content. The elastic modulus of WT streamers (*E* = 5.1 kPa ± 0.9 kPa) was double that of Pel-deficient streamers (*E* = 2.5 kPa ± 0.5 kPa) and 30% smaller than that of Pel-overproducer streamers (*E* = 7.2 kPa ± 0.9 kPa) (Fig. 3*A*, inset). The impact of Pel abundance on the effective viscosity was even more pronounced: WT streamers had approximately a 5-fold higher effective viscosity (*η* = 11.6 MPa s ± 1.7 MPa s) than Pel-deficient streamers (*η* = 2.7 MPa s ± 0.5 MPa s), while Pel-overproducing streamers had a viscosity 2.5-fold greater than WT streamers (*η* = 25.3 MPa s ± 2.9 MPa s) (Fig. 3*B*, inset). Therefore, since both *E* and *η* increased with increasing Pel abundance, we conclude that Pel causes stiffening in the viscoelastic response of biofilm streamer (Fig. 3).

**Figure 3.**
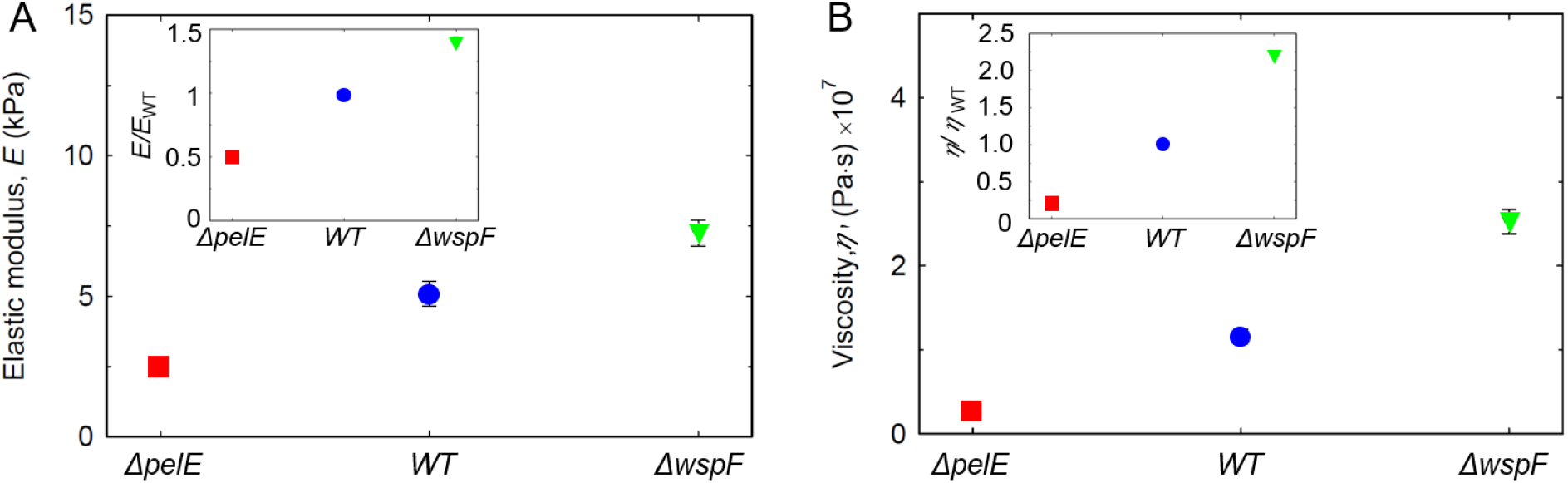
Pel stiffens biofilm streamers. (*A*) Elastic modulus, *E*, of biofilm streamers formed by *P. aeruginosa* PA14 *ΔpelE* (red), *WT* (blue) and *ΔwspF* (green) cells attached to a 50-μm pillar. Points show the average of the 5 independent average values associated to each bacterial batch The data were collected for 5 independent bacterial batches, each one prepared on a different day. The error bar is the standard deviation of the mean. Values were obtained by analyzing the deformation *in situ* of a portion of filament between 400 μm and 1 mm from the pillar. In the inset, the elastic modulus is rescaled by the elastic modulus of PA14 WT, *E/E*_WT_. (*B*) Effective viscosity, *η*, measured in the same experiments as *A*. In the inset, the effective viscosity is rescaled by the viscosity of PA14 WT, *η*/*η*_WT_. Error bar represent the standard error of the mean.

### eDNA degradation prevents streamer formation

DNA degradation caused by treatment with the enzyme DNase I completely prevented the formation of streamers by PA14 WT, demonstrating that eDNA is an essential structural component of streamers in PA14. PA14 WT did not form any streamers over 24 h when treated with DNase I solution, together with its activators CaCl2 and MgCl2, from the start of the experiment (Fig. 4*A, C, D*, Fig. S5*I*). DNase treatment also prevented the accumulation of biomass around the pillar (Fig. 4*C, E*). Phase-contrast imaging confirmed that streamers were not formed under DNase I treatment (Fig. S5*B, D, F, H*). Control experiments with only CaCl_2_ and MgCl_2_ revealed streamer formation (Fig. S5*A, C, E, G*), confirming that DNase I was responsible for preventing streamer formation. In contrast to its effect on streamers and on the pillar surface, DNase I treatment only modestly hampered biofilm presence on the channel’s bottom surface, reducing bacterial surface-coverage *C*_surf_ by about 50% (Fig. 4*F*): this supports the conclusion that eDNA is more important (in fact, essential) in the formation of streamers than in the formation of surface-attached biofilms. By focusing on early-stage biofilm formation (*t* < 24 h), we did not investigate the maturation of the biofilm, which according to the literature is prevented by DNase I (42). In the Pel-deficient case, DNase I also completely prevented streamer formation and biofilm accumulation around the pillar surface, but its effect on the channel’s surface was even weaker than for the WT (Fig. S6, Fig. 4*D–F*).

**Figure 4.**
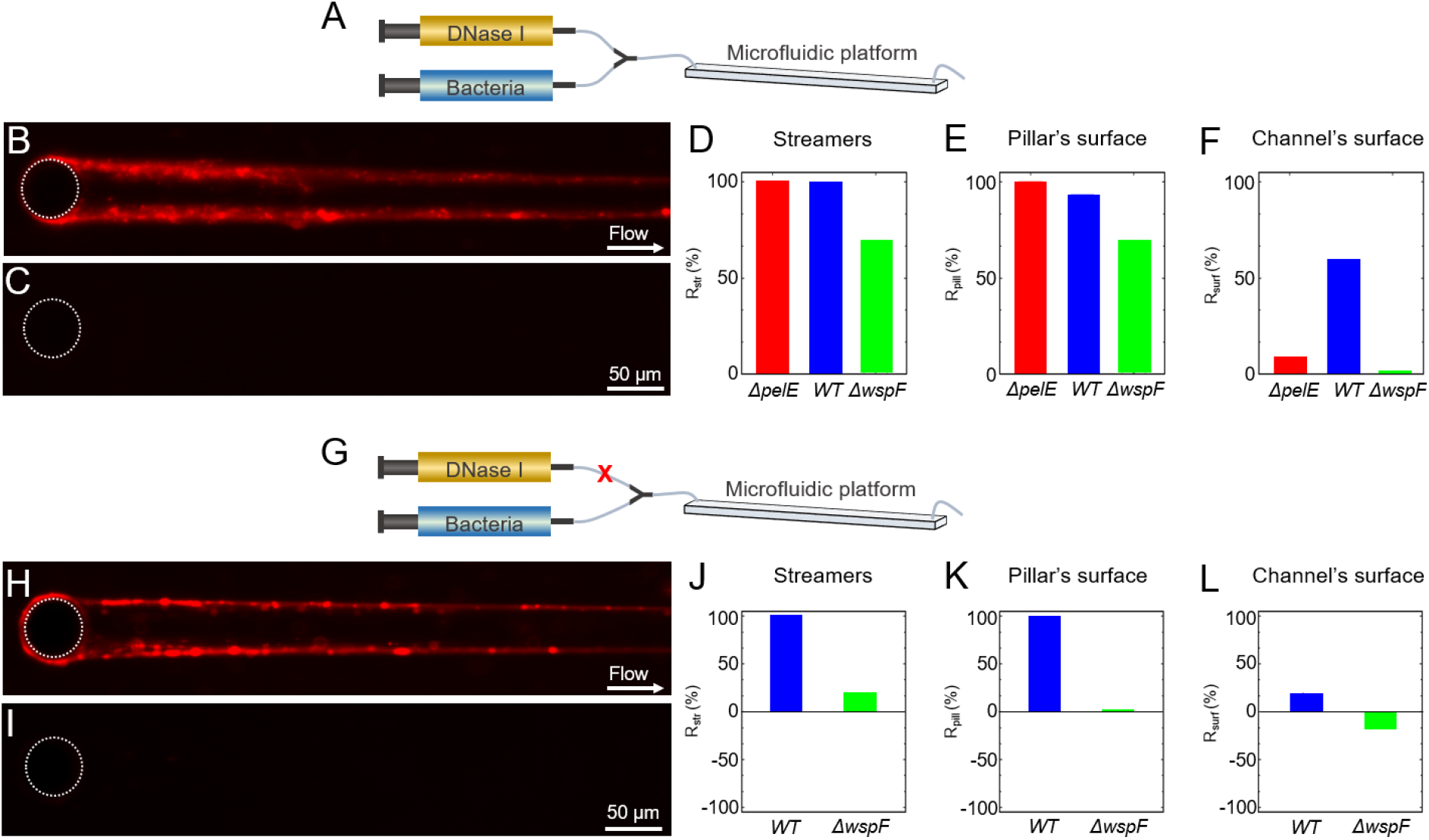
DNase I prevents biofilm streamer formation. (*A*) Schematic of the flow configuration during the experiment in which DNase I flow is injected at the beginning of the experiment. (*B,C*) Fluorescence images of biofilm streamers formed by *P. aeruginosa* PA14 WT cells attached to a 50-μm pillar after 24 h of continuous flow of a dilute bacterial suspension at *U* = 2 mm/s untreated (*B*) or treated (*C*) with 1 mg/mL DNase I. (*D*–*F*) Reduction in comparison to a control channel with no DNase treatment, *R*_str_, after 24 h of DNase I treatment calculated for the fluorescence intensity of the streamers, 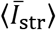, (*D*), *R*_pill_, for the fluorescence intensity of the biofilm around the pillar surface, 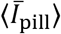, (*E*), and *R*_surf_, for the surface coverage, *C*_surf_, (*F*), for *P. aeruginosa* PA14 *ΔpelE* (red), WT (blue) and *ΔwspF* (green). (*G*) Schematic of the flow configuration during the experiment in which DNase I flow is started after 21 h post inoculation. The red cross represents a valve, which is kept close during the unperturbed streamers growth, and opened after 21 h when DNase I flow is started. (*H,I*) Fluorescence images of the biofilm streamers formed by *P. aeruginosa* PA14 WT cells attached to a 50-μm pillar after 21 h of continuous flow of a diluted bacterial suspension at *U* = 2 mm/s (*H*) and of the same streamer after 3 h of treatment with 1 mg/mL DNase I (*I*). (*J–L*) Reduction in comparison to a control channel with no DNase treatment, *R*_str_, of the signal after 3 h of DNase I treatment for the fluorescence intensity of the streamers, 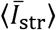, (*J*), *R*_pill_, for the fluorescence intensity of the biofilm around the pillar surface, 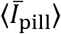, (*K*), and *R*_surf_, for the surface coverage, *C*_surf_ (*L*) for *P. aeruginosa* PA14 WT (blue) and *ΔwspF* (green).

An over-production of Pel in the biofilm formed by PA14 *ΔwspF* displayed a reduced effect of the DNase I treatment, represented by an approximately 60% reduction in streamer formation and colonization of the pillar surface (Fig. 4*D*, *F*). Additionally, the thickness of the biofilm on the pillar measured using phase-contrast microscopy seemed to increase during the treatment with DNase I and had an opposite trend as compared to the PI fluorescent signal (Fig. 4*E*, Fig. S7*A-E*). The biofilm structures that survive the DNase I treatment showed a high Pel content, as observed by staining Pel with WFL in the biofilm remaining after the treatment (Fig. S8*C, G*). eDNA is colocalized with Pel within these structures, as shown by staining (Fig. S8*D, H*). This suggests that Pel protects eDNA from the DNase I treatment. In line with the higher surface colonization ability of PA14 *ΔwspF* (Fig. 2*I*), surface coverage was not affected by DNAse I treatment (Fig. S7*G*, Fig. 4*F*), confirming that Pel plays a prominent role in the formation of a surface-attached biofilm.

To understand whether DNase I treatment could also effectively remove streamers after their formation, we exposed established streamers of PA14 WT and PA14 *ΔwspF* to DNase I. This treatment completely removed a 21 h PA14 WT streamer, along with the biomass on the pillars, within 30 min (Fig. 4*J, K*; Fig. S5*P, Q*), whereas it only reduced biofilm presence on the channel’s surface by 17% (Fig. 4*L*). Phase-contrast microscopy confirmed that DNase I treatment completely removed streamers (Fig. S5). In contrast, streamers formed by PA14 *ΔwspF* were reduced by 20% (Fig. 4*J*, Fig. S7*H*) and almost unaffected observing the Pel-rich biofilm formed around pillars (Fig. 4*K*; Fig. S7*O*). The biofilm on the channel’s surface was also unaffected and continued to increase during the treatment (Fig. 4*L*; Fig. S7*P*), showing a 15% increase during the 3 h treatment (Fig. 4*L*).

To explore and highlight the consequences of DNase I treatment on streamer formation in topographically more complex environments, we performed experiments with PA14 WT in a microfluidic model of a porous medium composed of thousands of identical pillars. When a dilute suspension of PA14 WT was flown through the porous medium, two different biofilm morphologies could be identified: in the first millimeter, a dense biofilm clog formed, whereas the downstream pillars supported a dense network of streamers, each akin to those observed on isolated pillars (Fig. S7*A*). DNase I treatment prevented clogging when started before streamers formation (Fig. S9) and reduced clogging when started after a mature streamers network had formed (Fig. S10). When DNase I was added to the flow from the beginning of the experiment, no streamers formed for the 23 hours of the experiment (Fig. S9, Movie S1). This observation further strengthens the conclusion that eDNA filaments are the structural elements of streamers, which in turn cause clogging. Thus, preventing the formation of streamers by removing eDNA could avoid or delay clogging. In experiments in which a 20 h WT streamers network was treated with DNase I, the efficacy of the treatment depended on the biofilm morphology. While eDNA in the clog was removed, resulting in shrinkage of the biofilm (Fig. S10*A*, *B*), some biomass remained, which most likely was composed of Pel. In contrast, the network of streamers was efficiently removed by DNase I (Fig. S10*C*, *D*), confirming that in PA14 WT, eDNA is essential for the integrity of streamers.

### Antibiotic-induced eDNA release promotes streamer formation

In further experiments, we observed that a higher abundance of eDNA resulted in thicker streamers. The eDNA concentration in the system can be increased by exposing bacteria to ciprofloxacin, an antibiotic that induces cell lysis and consequently the release of cytosolic contents, including DNA (57, 65). We performed experiments in which PA14 WT cells were exposed to three ciprofloxacin concentrations below the minimum inhibitory concentration (MIC), namely 0.005 μg/mL, 0.01 μg/mL and 0.02 μg/mL, as they were flown through the microfluidic device with isolated pillars (Materials and Methods; Fig. S11). We observed that increasing concentrations of ciprofloxacin caused an increase in the diameter of the streamer (Fig. 5*A–D*) and in the eDNA concentration in the streamer, measured as the average red fluorescent intensity, 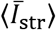 (Fig. S12*A*). The difference between the different ciprofloxacin concentrations in 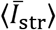 was initially small (*t* < 15 h; dotted line in Fig. S12*A*, red open triangles in Fig. 5*E*) and then rapidly increased (Fig. S12*A*). After 24 h, 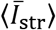 was tenfold higher in streamers exposed to 0.02 μg/mL of ciprofloxacin than in the untreated control (red filled triangles in Fig. 5*E*). 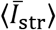 was linearly correlated with ciprofloxacin concentration at both 15 h and 24 h (Fig. 5*E*). The same trend is displayed by the concentration of single-stranded DNA (ss-DNA) measured in the effluent of the devices (Material and Methods; gray squares in Fig. 5*E*). Taken together, these data suggest that the increase in diameter of streamers is determined by an increased concentration of eDNA.

**Figure 5.**
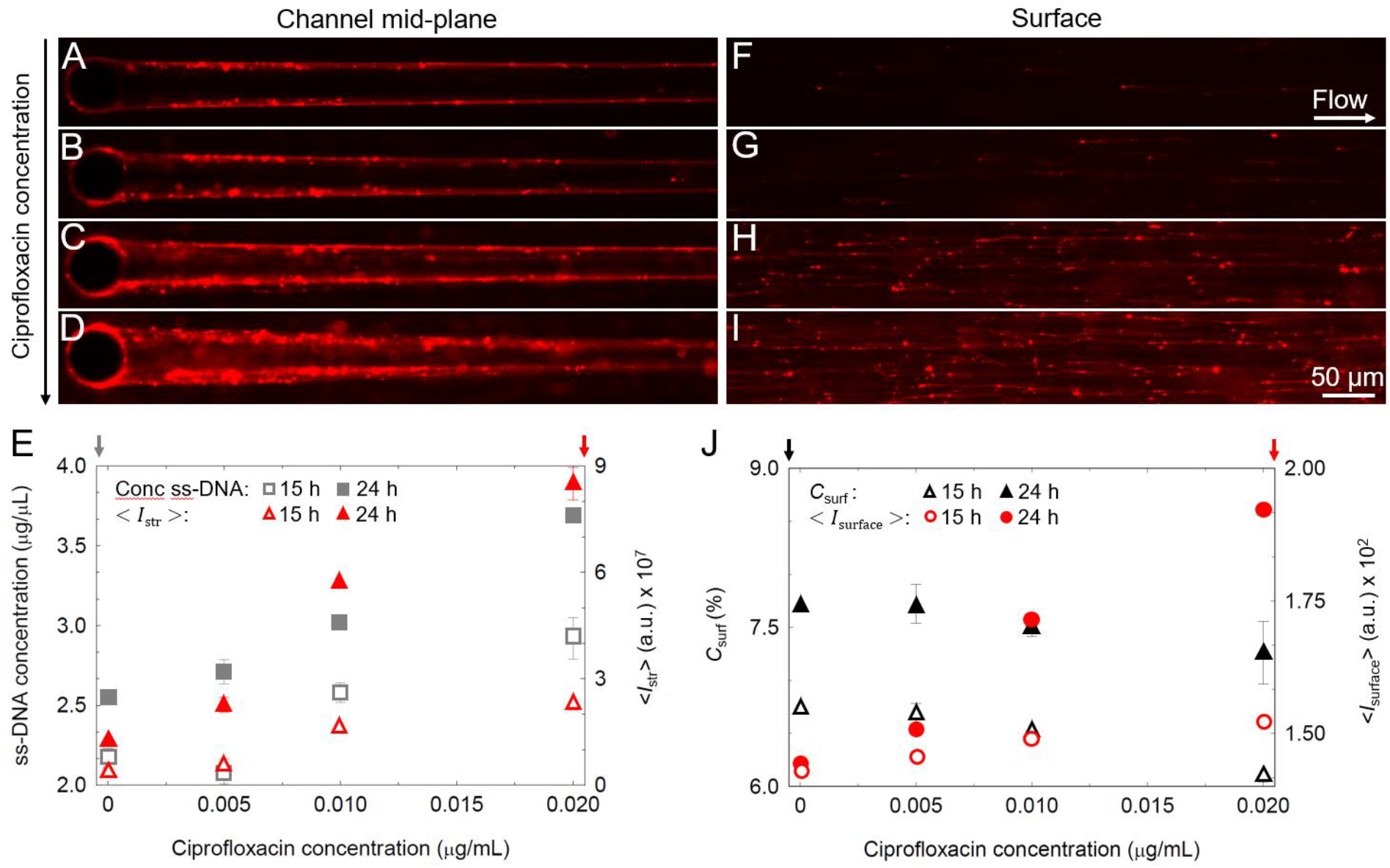
Ciprofloxacin increases biofilm streamer formation. (*A–D*) Fluorescence images of the biofilm streamers formed by *P. aeruginosa* PA14 WT attached to a 50-μm pillar after 24 h of continuous flow at *U* = 2 mm/s of a dilute bacterial suspension containing CPFLX at concentrations of 0 μg/mL (*A*), 0.005 μg/mL (*B*), 0.01 μg/mL (*C*) and 0.02 μg/mL (*D*). Images were taken at channel mid-depth. (*E*) Concentration of single-stranded DNA (ss-DNA) in solution (gray squares, left axis) and average red fluorescence intensity of the streamers, 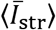, (red triangles, right axis) as a function of CPFLX concentration, measured after 15 h (open symbols) and 24 h (filled symbols) of continuous flow of a dilute PA14 WT suspension. (*F–I*) Fluorescence images of the eDNA network formed on the surface in the same channels as *A–D*, with CPFLX at concentrations of 0 μg/mL (*F*), 0.005 μg/mL (*G*), 0.01 μg/mL (*H*) and 0.02 μg/mL (*I*). Images were taken on the glass surface of the channel 3 mm upstream from the pillar. (*J*) Surface coverage (percentage), *C*_surf_, (black triangles, left axis) and red fluorescence intensity, 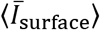, measured on the horizontal surface of the channel (red circles, right axis) as a function of CPFLX concentration, measured after 15 h (open symbols) and 24 h (filled symbols) in the same conditions as *A–D*.

Ciprofloxacin did not affect bacterial surface coverage, *C*_surf_, which showed similar values in all ciprofloxacin treatments (Fig. S12*B*, Fig. 5*J*), but it increased the abundance of eDNA on the surface (Fig. 5*F–I*), quantified by the average red fluorescent intensity of the surface, 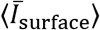 (red circles in Fig. 5*J*; Methods), increases linearly with the concentration of the antibiotic and therefore ss-DNA in solution. This observation strengthens the hypothesis that the increased growth of streamers is due to higher eDNA concentration in solution rather than cell capture, since the numbers of cells on the surface of the channel and in the effluent solution were comparable, whereas the concentration of eDNA in the effluent solution was dependent on the antibiotic concentration (Fig. 5*E*). A similar trend was observed with the Pel-deficient PA14 *ΔpelE* mutant (Fig. S13), which showed a small increase in streamer size with increasing ciprofloxacin concentration, and in the Pel-overproducing PA14 *ΔwspF* mutant (Fig. S13), which showed a more pronounced increase in streamer diameter. We thus speculate that the absence of Pel reduces the eDNA binding sites available on the streamers and consequently its absorption, thus reducing the size of the streamers. Similarly, Pel overproduction enhances eDNA absorption and streamer growth.

## Discussion

We have shown that eDNA is the fundamental structural component of *P. aeruginosa* biofilm streamers. Our observations demonstrated that the backbone of biofilm streamers consists of eDNA. The exopolysaccharide Pel is a further component of streamers, yet not an essential one, since a Pel-deficient mutant was still able to form streamers. In previous work (18, 19, 22), a Pel-deficient mutant did not form streamers in geometries where the filament had to cross the streamlines of the flow, as is the case in a channel with a series of corners. In contrast, our results in a geometry in which streamer growth occurred along the direction of streamlines show that a Pel-deficient mutant did form millimeter-long streamers. We hypothesize that the extensional component of the flow around the pillar may have favored the formation of streamers. Indeed, as previously shown using λ-phage DNA, a shear flow can induce and control the assembly of DNA, when the condition *γ τ* > 1 is satisfied, where *γ* is the flow shear rate and *τ* is the relaxation time of the DNA molecule (66, 67). In our configuration (Supplementary Information) we are indeed in the regime where the flow can favor DNA assembly and consequently the creation of streamers.

Our data indicate that the quantity of Pel secreted by bacteria affects the morphology of streamers. The Pel-deficient strain was found to have limited coverage of the flat surface, as reported in previous work (29, 37, 38), in which deficiency in Pel production was found to reduce the surface colonization ability and prevent the formation of a mature surface-attached biofilm. We found that Pel also promotes the formation of cell aggregates along the streamer and is mainly localized within these aggregates, whereas eDNA forms the millimeter-long continuum thread that supports the streamer structure. Interestingly, an increase in Pel corresponds to an increase of bacterial aggregates, supporting the idea that Pel promotes cell-cell adhesion not only in the surface-attached biofilm configuration (31), but also within suspended streamer filaments. Comparison of results using Pel-mutants showed that Pel has an important effect on the streamers’ rheological behavior: the higher the concentration of Pel, the higher the elastic modulus and effective viscosity of the streamers, and thus their stiffness.

eDNA is an abundant and structurally important component of *P. aeruginosa* surface-attached biofilms (41, 42). Evidence has emerged that eDNA in surface-attached biofilms is localized in distinct patterns that depend on the age of the biofilm and that this, in turn, determines the biofilm structure: young colonies present eDNA mostly on the upper surface, old colonies near the substratum (41). Here we have demonstrated that, for streamers, eDNA constitutes the skeleton of the streamer filaments. This observation is compatible with a previous report (19), where staining of streamers formed by *P. aeruginosa* PA01 with *Triticum vulgaris* (WGA) and *Canavalia ensiformis* (ConA) lectins allowed the visualization of the exopolysaccharide aggregates in the proximity of corners, but not of the EPS filaments spanning the channel, suggesting that the filaments were formed by eDNA.

In addition to our direct visualization of eDNA distribution within streamers (Fig. *2A-C*), the role of eDNA is further supported by our experiments with DNA degrading enzymes. In the Pel-deficient and wild-type strains, DNase I treatment completely prevented the formation of streamers, in both single-pillar (Fig. 4)) and porous media configurations (Fig. S9), and even established streamers. In Pel-overproducer streamers, treatment with DNase I was less effective and did not lead to complete streamers disruption (Fig. S7). When Pel is abundant in the filament, eDNA degradation leads only to a partial disruption of the biofilm matrix. This suggests that the Pel-eDNA ionic interaction shields the eDNA from the enzymatic action of DNase I or helps to maintain the degraded eDNA fragments within the biofilm matrix.

The key characteristic of eDNA in the context of biofilm formation is its binding affinity (34, 68). It is known that in mature mushroom structures formed by *P. aeruginosa* PA14, eDNA is found in the stalk where it binds to the positively charged exopolysaccharide Pel, resulting in increased structural stability (34). It has recently been shown that in surface-attached biofilms eDNA can be stabilized by proteins, resulting in the formation of lattice structures (68). Here, we showed that eDNA–Pel cross-links are not essential for the formation of biofilm streamers but significantly contribute to shaping their morphology and mechanical properties. In fact, we have shown that the Pel-overproducer strain generates shorter streamers with a larger diameter and large cell aggregates close to the pillar and that, conversely, the Pel-deficient strain forms long and slender filaments, easily shaped by the flow. Based on these observations, we hypothesize that the magnitude of the flow at which streamers are formed will have a significant impact on their morphology. Hence, the final morphology appears to be determined by the interplay between the force exerted by the flow and the biofilm matrix composition.

Using mutant strains in Pel production, we were also able to quantify the structural and mechanical properties of the eDNA-Pel interaction by directly measuring the elastic and viscous response to controlled mechanical stress. Interestingly, in our experiments, the ratio of elastic modulus and effective viscosity, which represents the elastic relaxation time, scales according to an almost universal relation previously proposed for biofilms (13) (Fig. S14). Moreover, we show that by causing an increase in both the elastic modulus and the viscosity, Pel enhances the stiffness of the streamer. Pel is a very small molecule (0.5 kDa) (27), even compared to fragments of eDNA (one base pair of double-stranded DNA is 0.65 kDa). Given that the opposite charge of eDNA and Pel may favor the creation of ionic bonds (27), we hypothesize that Pel molecules interconnect eDNA chains, thus reducing the slip between them and consequently affecting the response to deformations of the whole biofilm structure. These findings depict the streamers formed by *P. aeruginosa* PA14 as a bi-component network, formed by a combination of a short-chain (Pel) and a long-chain (eDNA) compound, connected by ionic interactions: networks with these characteristics, in which the shorter chain acts as an energy-dissipating element and improves the mechanical properties of the material, are known as double-network gels (69). The analogy between biofilm streamers and synthetic double-network gels – which have been recently developed to create hydrogels more resistant to deformations – suggests that the key element for biofilm mechanical resistance could reside in the interaction between the different components of EPS, with an important role played by eDNA.

Our results show that induction of lysis of *P. aeruginosa* PA14 by treatment with a sub-lethal concentration of ciprofloxacin promotes streamer formation by increasing the concentration of eDNA, leading to an increase of the streamers biomass. A recent study demonstrated that sublethal antibiotic concentrations promoted bacterial aggregation and resulted in elevated susceptibility to intestinal expulsion (70). Our experiments provide further and stronger evidence that, by increasing the amount of eDNA, some antibiotic treatments could enhance the formation of biofilms by promoting the occurrence of streamers which, given their spatial localization, eventually lead to clogging in medical devices such as stents or catheters and to the spreading of infection due to biofilm detachment. Moreover, since it is becoming clear that bacterial eDNA plays an important role in potentiating inflammation (71, 72), these findings are paving the way for future studies on the effects of antibiotic treatments on biofilm-forming pathogens and the host immune response.

## Materials and Methods

### Bacterial cultures

Experiments were performed using *P. aeruginosa* strain PA14 WT, Pel deletion mutants PA14 *ΔpelE* and Pel overproducer strain PA14 *ΔwspF*, kindly provided by the laboratory of Prof. Leo Eberl at the Department of Plant and Microbial Biology, University of Zürich (Switzerland). Single colonies were grown from frozen stocks on Luria broth (LB) agar plates at 37 °C for 24 h. *P. aeruginosa* suspensions were prepared by inoculating 3 mL Tryptone broth (TB; 10 g/L tryptone) with cells from a single colony and incubating for 3 h at 37 °C, while shaking at 200 rpm. The suspensions were then diluted in fresh Tryptone Broth to final optical density OD_600_ = 0.01.

eDNA and Pel quantities were visualized and measured using fluorescence staining methods: for eDNA visualization, propidium iodide, PI (Sigma Aldrich) was added to the medium to a final concentration of 2 μg/mL, and for Pel visualization, Wisteria Floribunda, WFL (bioWorld) was dissolved in PBS buffer to a final concentration of 50 μg/mL. For experiments involving degradation of eDNA, DNase I (Sigma Aldrich) was dissolved in Tryptone broth to a final concentration of 1 mg/mL and CaCl_2_ and MgCl_2_ (final concentrations 0.12 mM) were added as activators. For antibiotic treatment experiments, ciprofloxacin (Sigma Aldrich) was first dissolved in 0.1 N HCl to a final concentration of 20 mg/mL and then further diluted in Tryptone broth to obtain solutions of final concentration 0.005 μg/mL, 0.01 μg/mL and 0.02 μg/mL. All samples of the antibiotic treatment experiment were assessed by high sensitivity double-stranded DNA assay and high sensitivity single-stranded DNA assay according to established protocols using Quibit 3.0 (ThermoFisher Scientific).

### Microfluidic assays

To analyze streamer formation in flow, we fabricated a polydimethylsiloxane (PDMS) microfluidic device with four channels on the same chip, each containing six cylindrical pillars of diameter 50 μm. The channel was 40 μm high and 1 mm wide. Pillars were located at the center of the channel and the distance between pillars was 5 mm. The flow was driven by a syringe pump (neMESYS 290N, CETONI, Germany). Prior to use, all microfluidic channels were washed with 2 mL of TB medium. A diluted PA14 bacteria suspension (OD_600_ = 0.01; cell concentration < 10^6^ cells/mL) was flown for 24 h. All experiments were performed at room temperature (T = 20 ± 0.5 °C). In the experiments in which DNase I was used, the DNase I solution was flown in a Y connector (P-514, IDEX) located before the inlet to avoid contact between the cells and the enzyme before the channel inlet (Fig. 4*A*, *G*). To perform DNase treatment on mature streamers, a shut-off valve (P-782, IDEX) was inserted between the syringe containing the DNase I solution and the Y connector (red cross in Fig. 3*G*). The shut-off valve was kept closed during streamer growth and then opened to expose the mature streamer to DNase I treatment. In the experiments in which ciprofloxacin was used, the antibiotic solution was flown in a Y connector (P-514, IDEX) located before the inlet to avoid contact between the cells and ciprofloxacin before the channel inlet, using the same configuration described for the DNase I treatments (Fig. 4*A*). For the experiment in which the model porous medium containing 75-μm pillars, the device was fabricated using polydimethylsiloxane (PDMS) and flow was driven by a syringe pump (neMESYS 290N, CETONI).

### Cell imaging and tracking

All imaging was performed on an inverted microscope (Ti-Eclipse, Nikon, Japan) using a digital camera (ORCA-Flash4.0 V3 Digital CMOS camera, Hamamatsu Photonics, Japan). Bacterial cells were imaged using phase-contrast microscopy (20× magnification). Biofilm composition was quantified using epifluorescence microscopy (20× magnification). During biofilm streamer growth, images were taken every 15 min both in phase contrast and epifluorescence, unless specified otherwise in the figure captions. During mechanical tests on biofilm streamers, images were taken before the tests in epifluorescence and once per second in phase-contrast during the tests. All images of biofilm streamers were obtained at channel mid-depth, while surface coverage was evaluated on the glass wall of the microfluidic channel in a region located 3 mm upstream of the pillars.

### Statistics and derivations

All image analysis was performed in Fiji-Image J (73). All images of streamers are examples from experiments that were repeated three times with consistent results.

The average fluorescence intensity of the streamers in each channel was calculated on the red fluorescence signal as

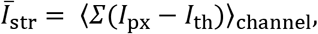

where *I*_px_ is the intensity of the pixels, and *I*_th_ is a threshold calculated as 1.2 times the average intensity of the background, *I*_back_, measured in the region of the channel on the side of the streamer. The sum was performed on a 550 μm × 100 μm area located downstream of each pillar (white dotted line in Fig. S3) and averaged over the six pillars contained in each channel. During each experiment, multiple channels with the same conditions were present and the average fluorescent intensity of the streamers 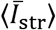 reported in the figures was obtained as an average over channels under the same experimental conditions. The same procedure was used to calculate the average fluorescence intensity on the pillar surface, 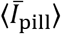. In this case, the sum was performed on an area of dimensions 55 μm × 100 μm located around the pillar (yellow dotted line in Fig. S3).

The diameter and length of streamers were measured on the fluorescence images of the red signal. The streamer was identified as the region of the image where the intensity of the pixels, *I*_px_, exceeded the threshold, *I*_th_. The diameter of the streamer was defined as the width of the filament in the direction perpendicular to the flow. The diameter of the streamer in the region close to the pillar, *d*_150_, was averaged over a distance of 150 μm from the pillar. The diameter of the streamer in the region downstream from the pillar, *d*_400_, was taken as the average of the width at a point located 400 μm from the pillar and a point located 1000 μm from the pillar. To measure the length of the streamer, *L*, images were stitched together using the Stitching Plugin in Fiji-Image J.

The coverage of the horizontal surface of the channel, *C*_surf_, was defined as the percentage of the surface covered by cells and was calculated on phase-contrast images of the surface of the channel. One region with dimensions 670 μm × 670 μm, located 3 mm upstream from a pillar and free from the presence of the streamer, was considered for each channel. Bacterial cells appeared as darker spots and their area was estimated as the number of pixels with an intensity lower than 0.8 × *I*_surface, ph_, where *I*_surface, ph_ is the average gray intensity of the bacteria-free surface acquired in phase-contrast. In Fig. 5, we report the average red fluorescence intensity of the surface, 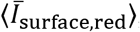, which was defined as the average red fluorescence intensity (corresponding to the PI signal) in the region with dimensions 670 μm × 670 μm used to calculate the surface coverage.

Values of *d*_150_, *d*_400_, 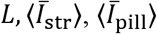 and *C*_surf_ reported in Fig. 2 were measured for each experiment and the values were averaged over two channels with identical experimental conditions. The error bars represent the standards deviations of the mean. The *Pel intensity* shown in the inset of Fig. 2*F* was calculated using the same definition of 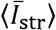 on single images of streamers formed after 20 h of continuous flow and stained first with WF for 30 min and then with PI for 30 min (sample images shown as Fig. S1 and S2). In this case, each value is obtained as an average over six pillars and experiments were repeated twice.

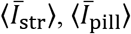 and *C*_surf_ reported in Fig. S5*I-K* were measured during the same experiment and the values are averaged over two channels. The same applies to Fig. S5*P-R*. In Fig. 4, the Reduction reported in panels *D-F* is defined as

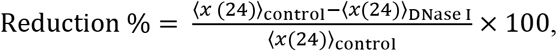

where *x(*24) represents the quantity reported in the main panel (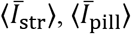 and *C*_surf_) and measured after 24 h. We refer to control as the channel in which no DNase I treatment was performed.

Reduction reported in Fig 4 panels *J-L* is defined as

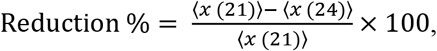

where *x(21)* represents the quantity reported in the main panel (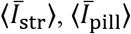 and *C*_surf_) and measured after 21 h of flow of Tryptone broth (before DNase I treatment) and *x(24)* the same quantity measured after 3 h of DNase I treatment. In this case, a negative reduction indicates an increase of the quantity during the DNase I treatment.

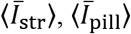 and *C*_surf_ reported in Fig. 5 were measured during the same experiment and the values are averaged over two channels. The single-stranded DNA concentration was measured in the effluent suspension of each channel, for the point at 15 h (corresponding to the liquid sampled in the period 14.5–15.5 h) and the point at 24 h (sampled 23.5–24.5 h).

The data reported in the Supplementary are measured during the same experiments and the values are averaged over two channels. All the experiments reported were performed at least three times and in each experiment a consistent trend was found.

### Mechanical tests

Rheological characterization was performed on streamers grown at a flow rate *Q* = 0.3 ml/h (average flow velocity 2 mm/s). *ΔwspF* streamers were tested after 14-15 h from the beginning of the experiment, while WT and *ΔpelE* after 20-22 h. Stress tests were performed by increasing the flow rate to 0.6 ml/h (average flow velocity 4 mm/s) for 5 min (red curve, Fig. S4*A*). While at this flow rate, the deformation of a portion of the streamer in the region between 400 μm and 1 mm from the pillar was imaged in phase contrast at 1 fps. Deformation was quantified by tracking the relative displacement of cell aggregates within the filaments, using the software Blender (https://www.blender.org/) (blue curve, Fig. S4*A*). The stress step applied in each experimental replicate was quantified using finite element code (COMSOL Multiphysics). The 3D model of the streamers was built for each replicate based on the eDNA signal of epifluorescence images acquired before each stress test. The results reported in Fig. 3 were calculated as the average of five independent bacterial batches, each one prepared on a different day (Fig. S4*B*, *C*), and the values of *E* and *η* for a single batch were an average over identical experiments. The error bar is the standard deviation of the mean.

## Acknowledgments

The authors acknowledge support from SNSF PRIMA grant 179834 (to E.S.), from Gordon and Betty Moore Foundation Marine Microbial Initiative Investigator Award GBMF3783 (to R.S.), and from Simons Foundation Grant 542395 (to R.S.) as part of the Principles of Microbial Ecosystems Collaborative (PriME).

## Supplementary Information

### eDNA assembly in flow

According to the literature about the aggregation of DNA molecules in shear flow (1, 2), shear flow can induce and control the assembly of λ-phage DNA when the condition *γτ* > 1 is satisfied, where *γ* is the flow shear rate and *τ* is the relaxation time of the molecule, which depends on the size of the DNA fragments. Under this condition, DNA molecules are stretched and deformed by flow, which exposes their ends, and their intermolecular interaction probability is increased, leading to inter-chain bonding and the formation of 3D elongated structures (1).

The shear rate values in a 5 μm-thick annulus around the pillar in the midplane of the channel, where eDNA aggregation is likely to happen, are in the range 280 *s*^−1^ < *γ* < 700 *s*^−1^.

The bulk values of the relaxation time, τ, can be estimated using

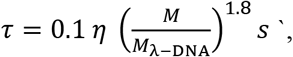

where *η* is the viscosity of the buffer (in our case water, *η* = 1 cP), *M* is the molecular weight of the eDNA molecule and *M_λ−DNA_* is the molecular weight of of λ-DNA (*M_λ−DNA_* = 48.5 kbp) (3).

Given the limiting values for γ, we can calculate the molecular weight *M* of the eDNA molecules required to satisfy the condition *γτ* > 1 as

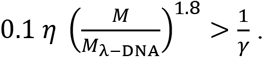

For *γ* =280 *s*^−1^, we obtain *M* > 7.6 kbp, while for *γ* =700 *s*^−1^, *M* > 4.6 kbp. In conclusion, in our configuration, the assembly of eDNA molecules with *M* > 4.6 kbp is promoted by the flow.

Since eDNA released by cell lysis is similar to chromosomal DNA (4, 5) and the genome of *P. aeruginosa* is 6.3 Mbp (6), the fragments present are likely of a size for which assembly is promoted by the flow.

**Fig. S1.**
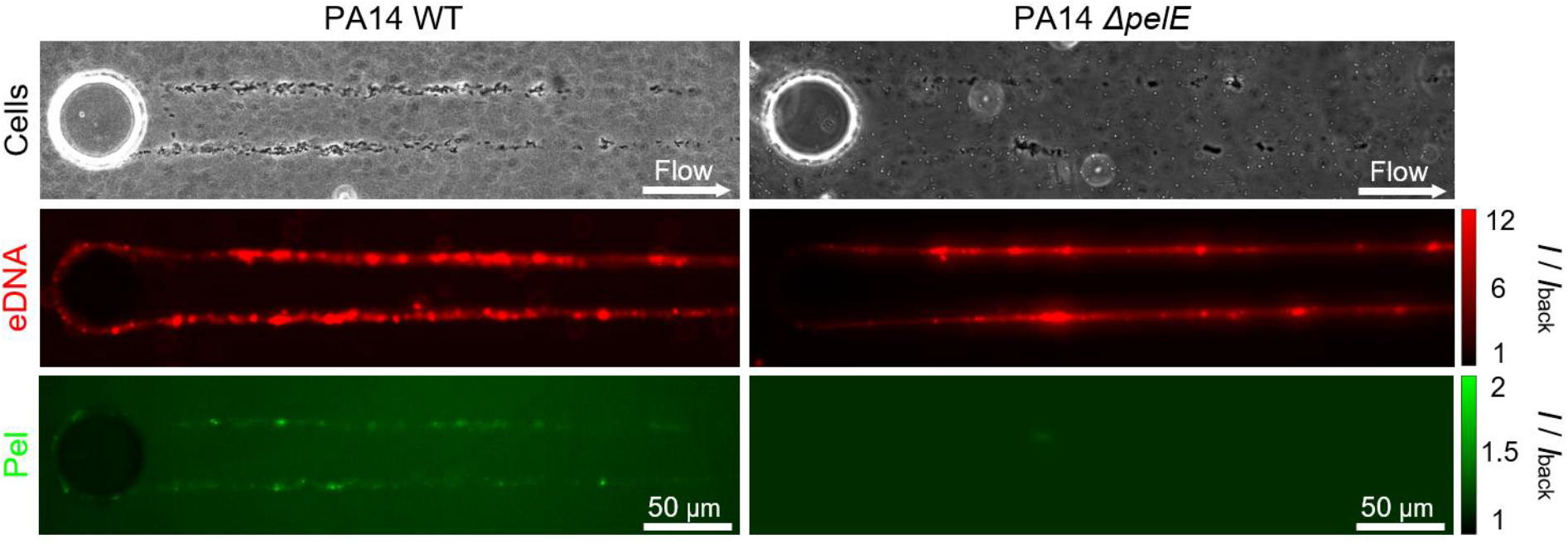
Streamer formation by the *P. aeruginosa* PA14 wild type and the Pel-deficient Δ*pelE* strain. Phase-contrast and fluorescence images of the biofilm streamers formed by *P. aeruginosa* PA14 WT (left) and Δ*pelE* (right) cells attached to a 50-μm pillar after 20 h of continuous flow of a dilute bacterial suspension at *U* = 2 mm/s. The images were taken at channel mid-depth. Bacterial cells were imaged in phase contrast (top), eDNA was stained using red-fluorescent PI (2 μg/mL; middle) and Pel was stained using the green-fluorescent WFL (50 μg/mL; bottom). To ease the comparison of the fluorescence images, the intensity was divided by the average fluorescence intensity in the region of the channel not occupied by the streamer.

**Fig. S2.**
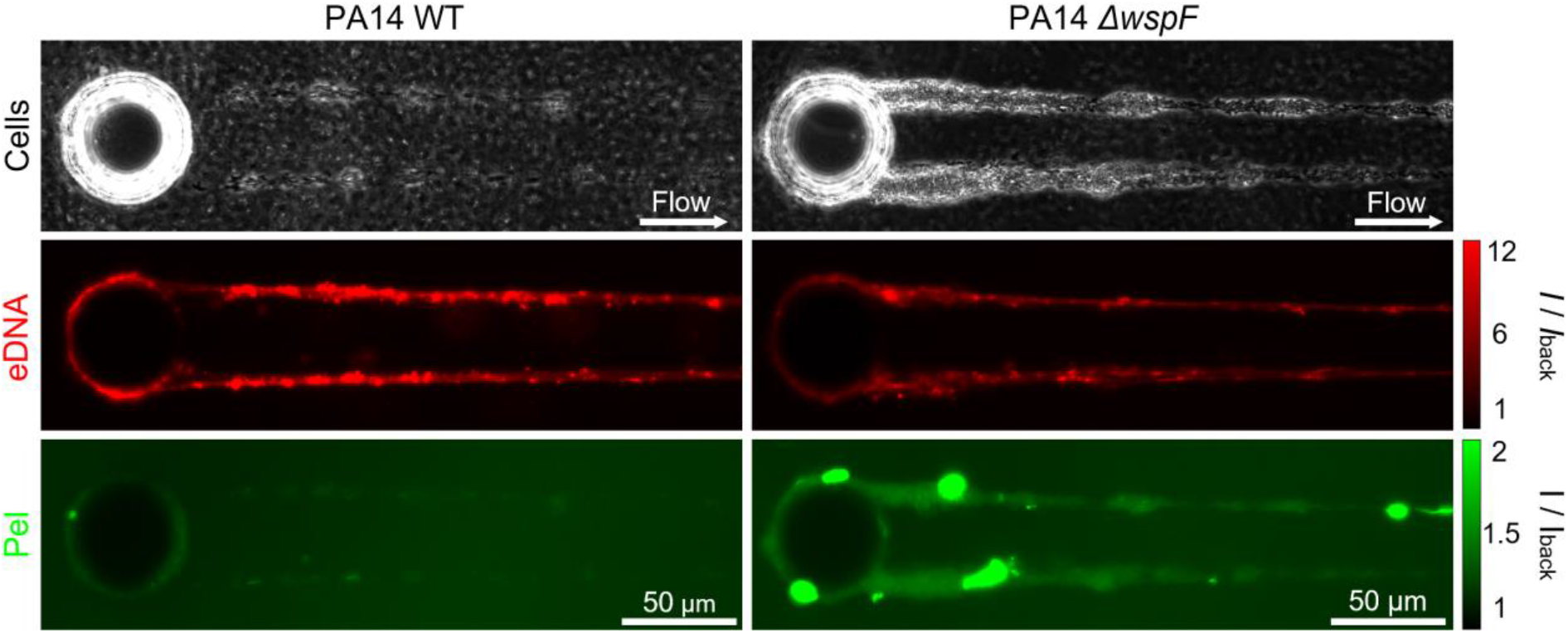
Streamer formation by *P. aeruginosa* PA14 WT and the *Pel* overproducer *ΔwspF* strain. Phase-contrast and fluorescence images of the biofilm streamers formed by *P. aeruginosa* PA14 WT (left) and *ΔwspF* (right) cells attached to a 50-μm pillar after 20 h of continuous flow of a dilute bacterial suspension at *U* = 2 mm/s. The images were taken at channel mid-depth. Bacterial cells were imaged in phase contrast (top), eDNA was stained using red-fluorescent PI (2 μg/mL; middle) and Pel was stained using the green-fluorescent WFL (50 μg/mL; bottom). To ease the comparison of the fluorescence images, the intensity was divided by the average fluorescence intensity in the region of the channel not occupied by the streamer.

**Fig. S3.**
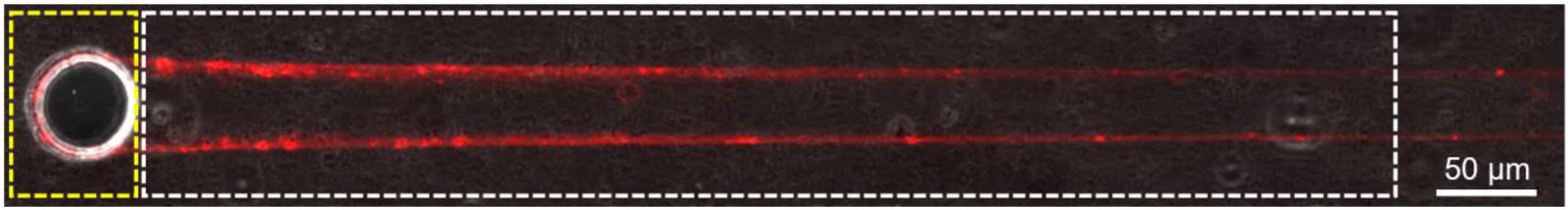
Sample regions selected to quantify streamer fluorescence intensity. Average fluorescence intensity of the streamers and of the biofilm around the pillar surface were measured in the regions of the image contained within the 550 μm × 100 μm dashed white box and the 55 μm × 100 μm dashed yellow box, respectively.

**Fig. S4.**
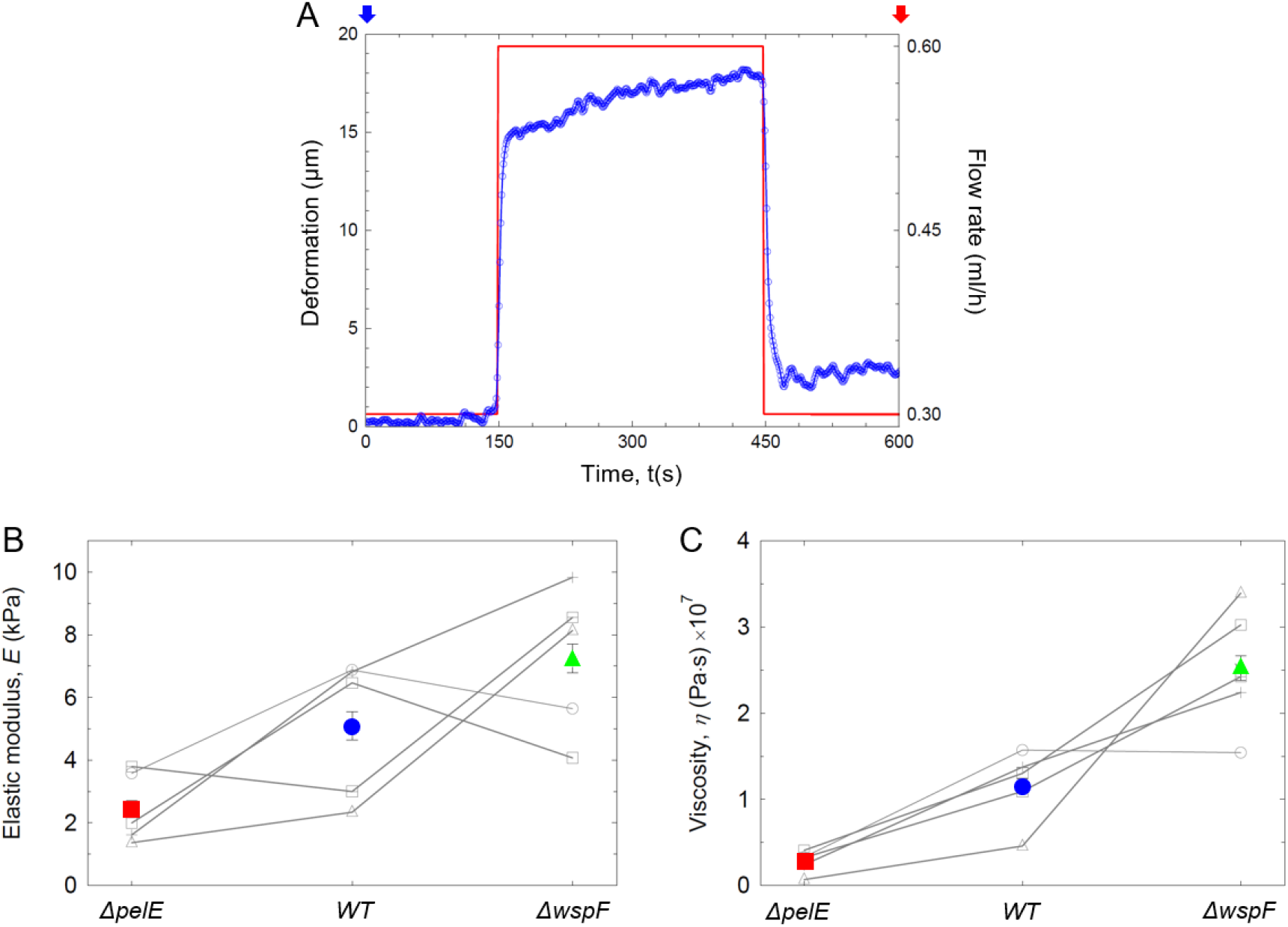
Mechanical test of biofilm streamer rheological properties. (*A*) Deformation of a PA14 WT streamer (blue curve, left axis) undergoing the mechanical tests. The streamer was deformed by increasing the flow rate (red curve, right axis) from 0.3 ml/h (corresponding to *U* = 2 mm/s) to 0.6 ml/h (corresponding to *U* = 4 mm/s). (*B*) Elastic modulus, *E*, of the biofilm streamers reported in Fig. 3, obtained as the mean of five independent experimental replicates carried out on different days (gray symbols). (*C*) Viscosity, *η*, measured in the same experiments as Fig. 3, was obtained as the mean of five independent experimental replicates carried out on different days (gray symbols). The error bar is calculated as the standard error of the mean.

**Fig. S5.**
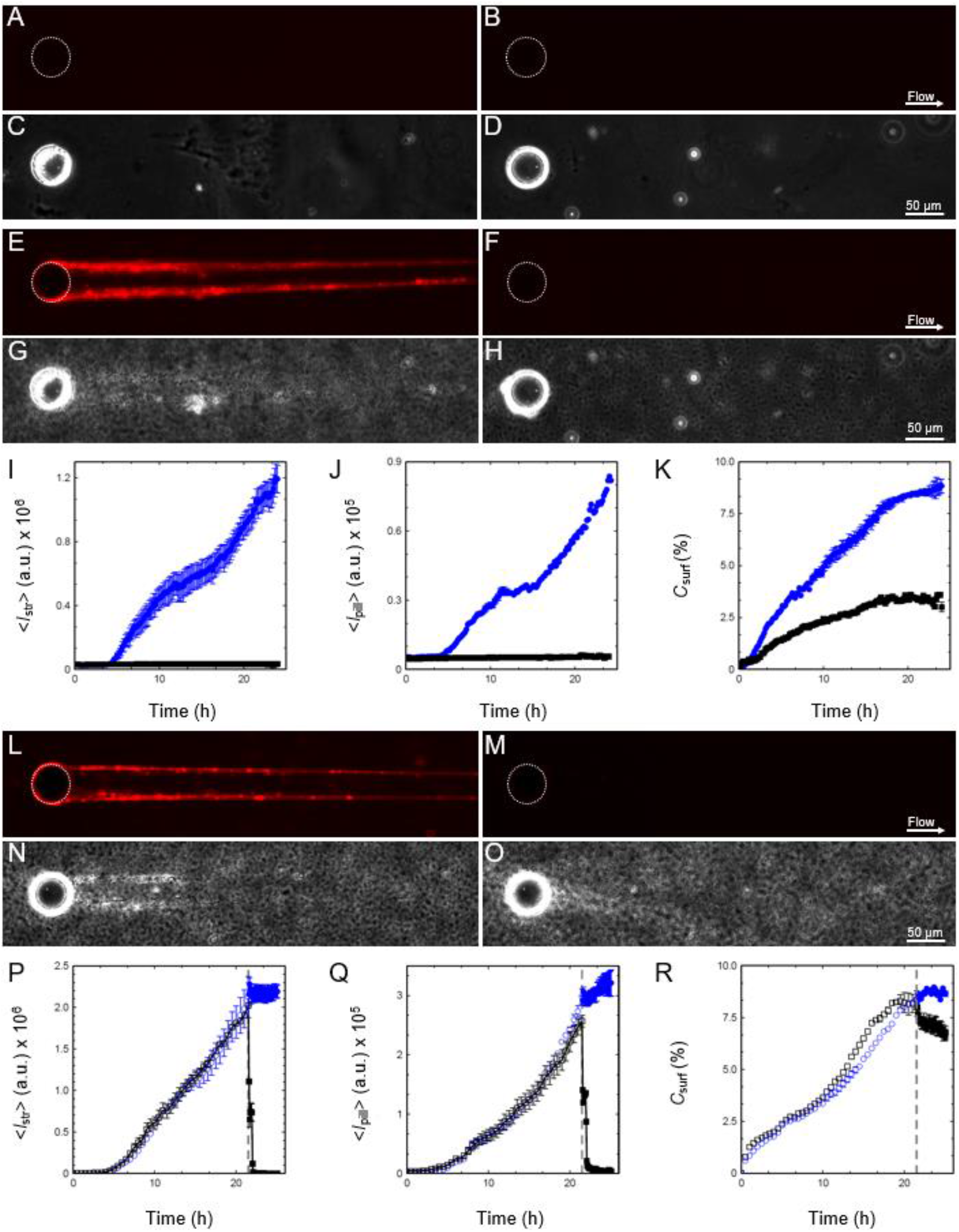
DNase I prevents streamer formation by PA14 WT and causes degradation of established streamers. (*A–H*) Fluorescence (*A, B, E, F*) and phase-contrast (*C, D, G, H*) images of the biofilm streamers formed by *P. aeruginosa* PA14 WT cells attached to a 50-μm pillar when the flow is started (*A-D*) and after 24 h of continuous flow (*E-H*) of a dilute bacterial suspension at *U* = 2 mm/s either untreated (*A, C, E, G*) or treated with 1 mg/mL DNase I (*B, D, F, H*). (*I-K*) Fluorescence intensity of the streamers, 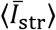 (*I*) and of the biofilm around the pillar surface, 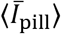 (*J*), and surface coverage, *C*_surf_ (*K*) as a function of time for the same suspension in *A* with no DNase treatment (blue circles) and *B* treated with DNase (black squares). Points show the mean and standard error of the mean of two replicates. (*L-O*) Fluorescence (*L, M*) and phase-contrast (*N, O*) images of the biofilm streamers formed by *P. aeruginosa* PA14 WT cells attached to a 50-μm pillar after 21 h of continuous flow of a diluted bacterial suspension at *U* = 2 mm/s (*L, N*) and of the same streamer after 3h of treatment with 1 mg/mL DNase I (*M, O*). (*P-R*) Fluorescence intensity of the streamers, 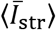 (*P*) and of the biofilm around the pillar surface, 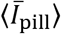 (*Q*), and surface coverage, *C*_surf_ (*R*) as a function of time, measured during 21 h of continuous flow of a diluted bacterial suspension in Tryptone broth (open symbols) and during the following 3 h during which streamers were exposed to flow of 1 mg/mL DNase I in Phosphate Buffer Saline, PBS, (black filled squares) or just PBS (blue filled circles). Points show the mean and standard error of the mean of two replicates.

**Fig. S6.**
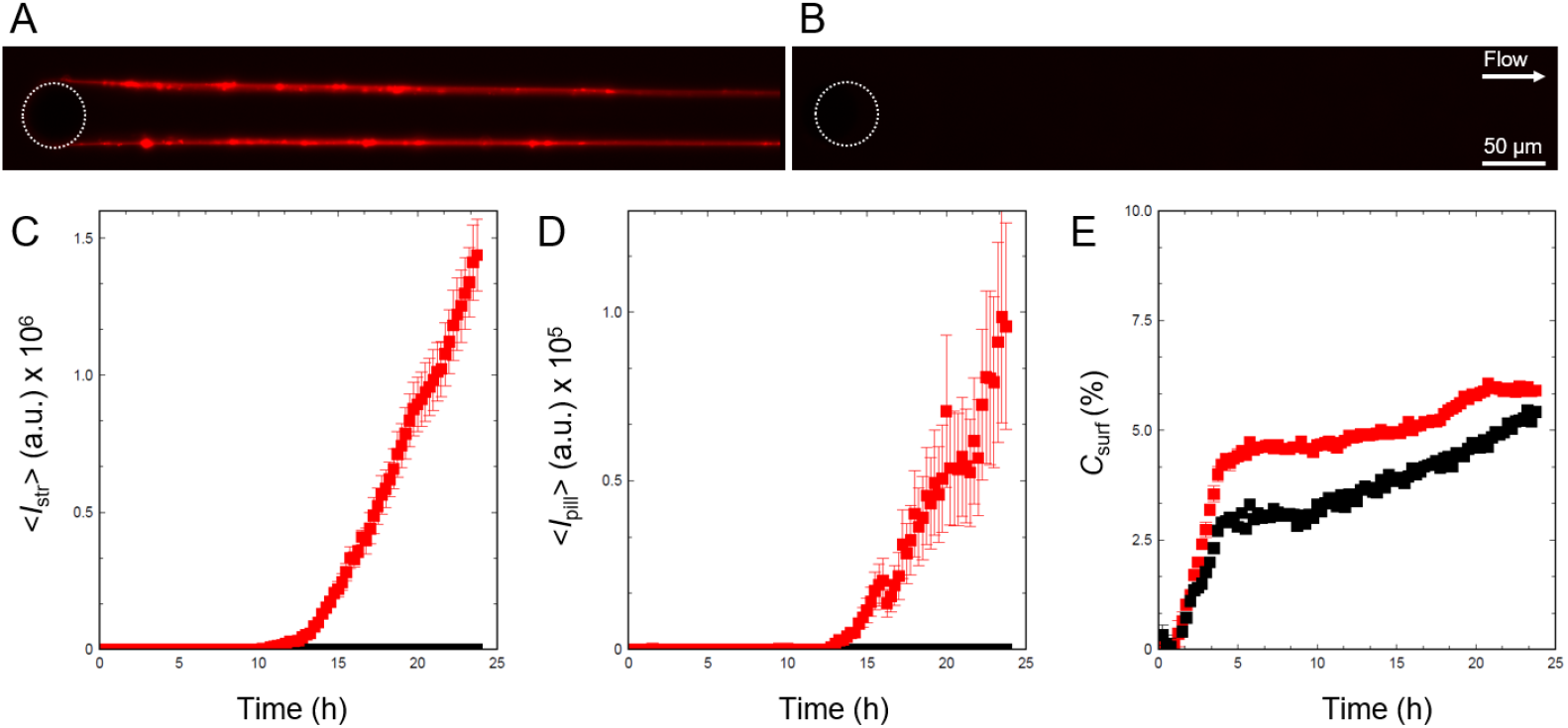
DNase I prevents streamer formation by PA14 *ΔpelE*. (*A-B*) Fluorescent image of the biofilm streamers formed by *P. aeruginosa* PA14 *ΔpelE* cells attached to a 50-μm pillar after 24 h of continuous flow of a diluted bacterial suspension at *U* = 2 mm/s untreated (*A*) and treated with 1 mg/mL DNase I (*B*). (*C-E*) Fluorescent intensity of the streamers, 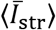 (*C*) and of the biofilm around the pillar surface, 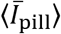 (*D*), and surface coverage, *C*_surf_ (*E*) as a function time for the same suspension in *A* (red squares) and *B* (black squares). Points show the mean and standard error of the mean of two replicates.

**Fig. S7.**
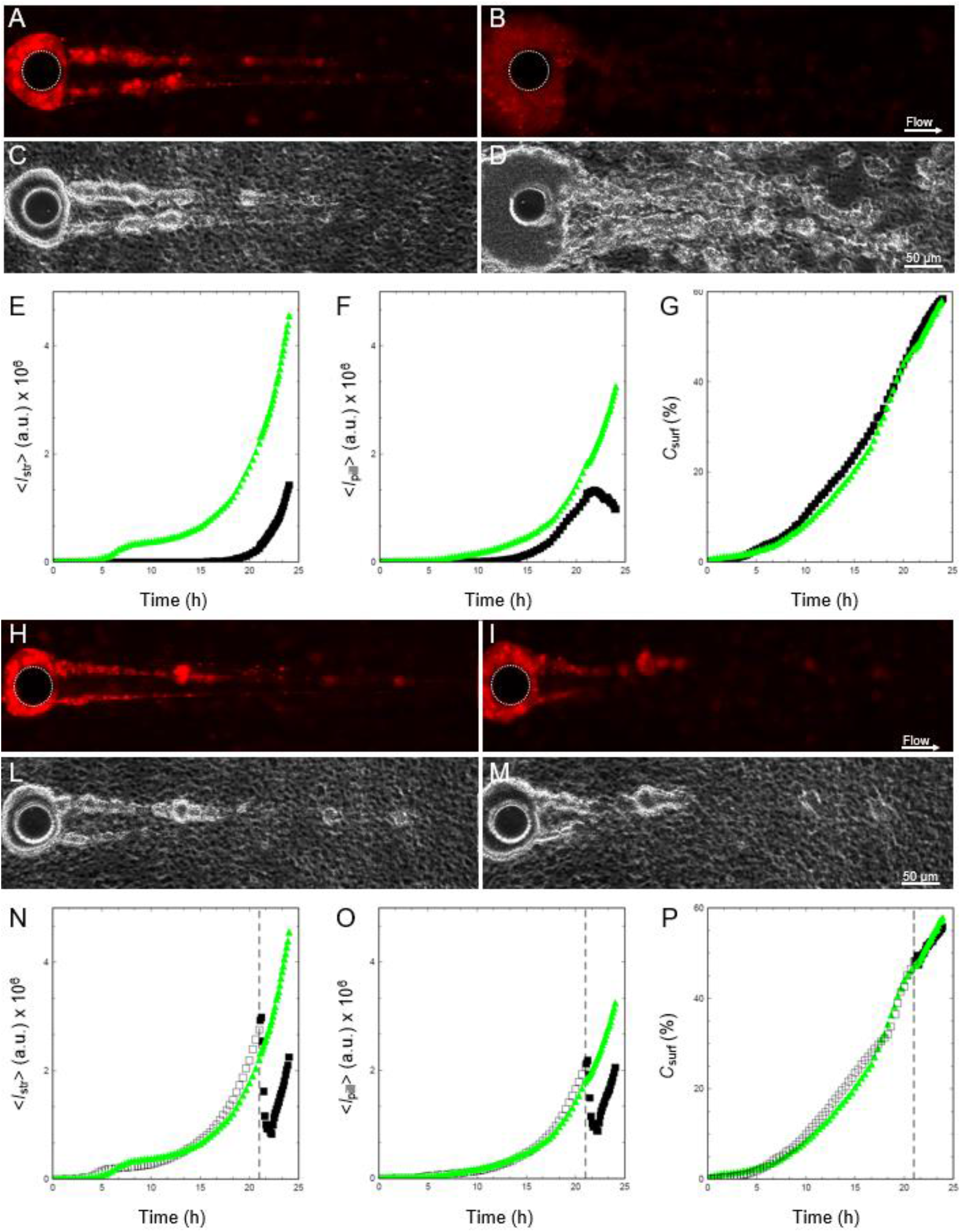
DNase I prevents streamer formation by PA14 *ΔwspF* and causes degradation of established streamers. (*A-D*) Fluorescent (*A, B*) and phase contrast images of the biofilm streamers formed by *P. aeruginosa* PA14 *ΔwspF* cells attached to a 50-μm pillar after 24 h of continuous flow of a diluted bacterial suspension at *U* = 2 mm/s untreated (*A, C*) and treated (*B, D*) with 1 mg/mL DNase I. (*E-G*) Fluorescence intensity of the streamers, 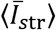 (*E*) and of the biofilm around the pillar surface, 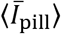 (*F*), and surface coverage, *C*_surf_ (*G*) as a function time for the same suspension in *A* (green triangles) and *B* (black squares). (*H-M*) Fluorescence (*H, I*) and phase contrast (*L, M*) images of the biofilm streamers formed by *P. aeruginosa* PA14 *ΔwspF* cells attached to a 50-μm pillar after 21 h of continuous flow of a diluted bacterial suspension at *U* = 2 mm/s (*H, L*) and of the same streamer after 3h treatment with 1 mg/mL DNase I (*I, M*). (*N-P*) Fluorescence intensity of the streamers, 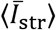 (*N*) and of the biofilm around the pillar surface, 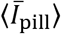 (*O*), and surface coverage, *C*_surf_ (*P*) as a function time measured during 21 h of continuous flow of a diluted bacterial suspension in Tryptone broth (open symbols) and during the following 3 h-treatment performed flowing 1 mg/mL DNase I in PBS (black filled squares) or just PBS (green filled triangles) for the same suspension in *H* (green triangles) and *I* (black squares).

**Fig. S8.**
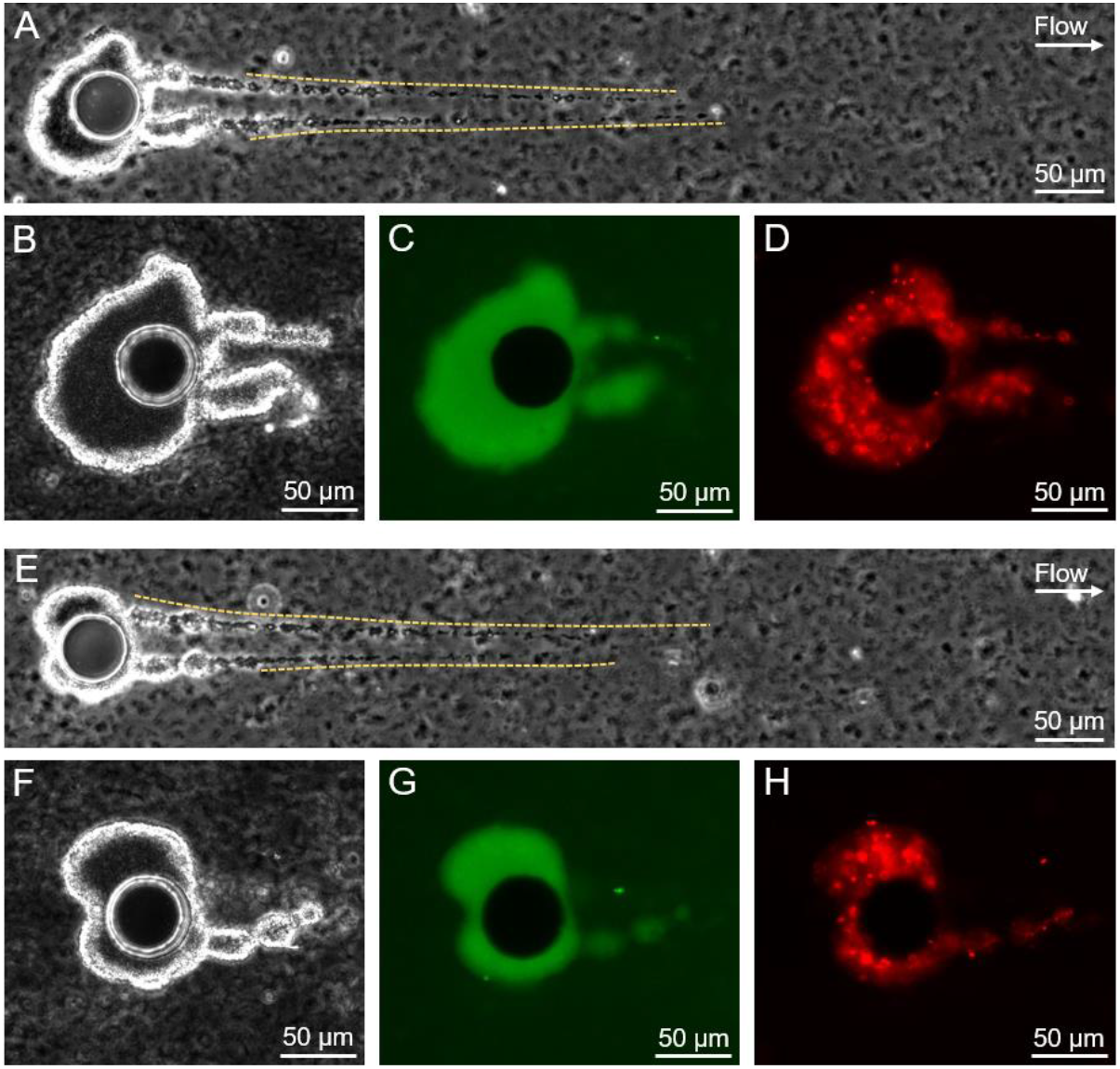
Pel reduces the effectiveness of DNase I treatment on PA14 *ΔwspF* streamers. (*A-H*) Phase contrast (*A, B, E, F*) and fluorescent (*C, D, G, H*) images of the biofilm streamers formed by *P. aeruginosa* PA14 *ΔwspF* cells attached to a 50-μm pillar after 21 h of a continuous flow of a dilute bacterial suspension at *U* = 2 mm/s (*A, E*) and of the same streamer after 3h treatment with 1 mg/mL DNase I (*B-D, F-H*). Yellow dotted lines in *A* and *E* indicate the portions of the streamer removed during the treatments. Bacterial cells are imaged in phase contrast, eDNA is stained using red-fluorescent PI (2 μg/mL; *D, H*) and Pel is stained using the green-fluorescent WFL (50 μg/mL; *C, G*). DNase I treatment destroys the filaments, as we can see comparing the images taken before (*A, E*) and after (*B, F*) the treatment. However, the Pel-containing aggregates (*C, G*) are not affected by the treatment and the Pel network inhibits the DNase I destructive action con eDNA (*D, H*).

**Fig. S9.**
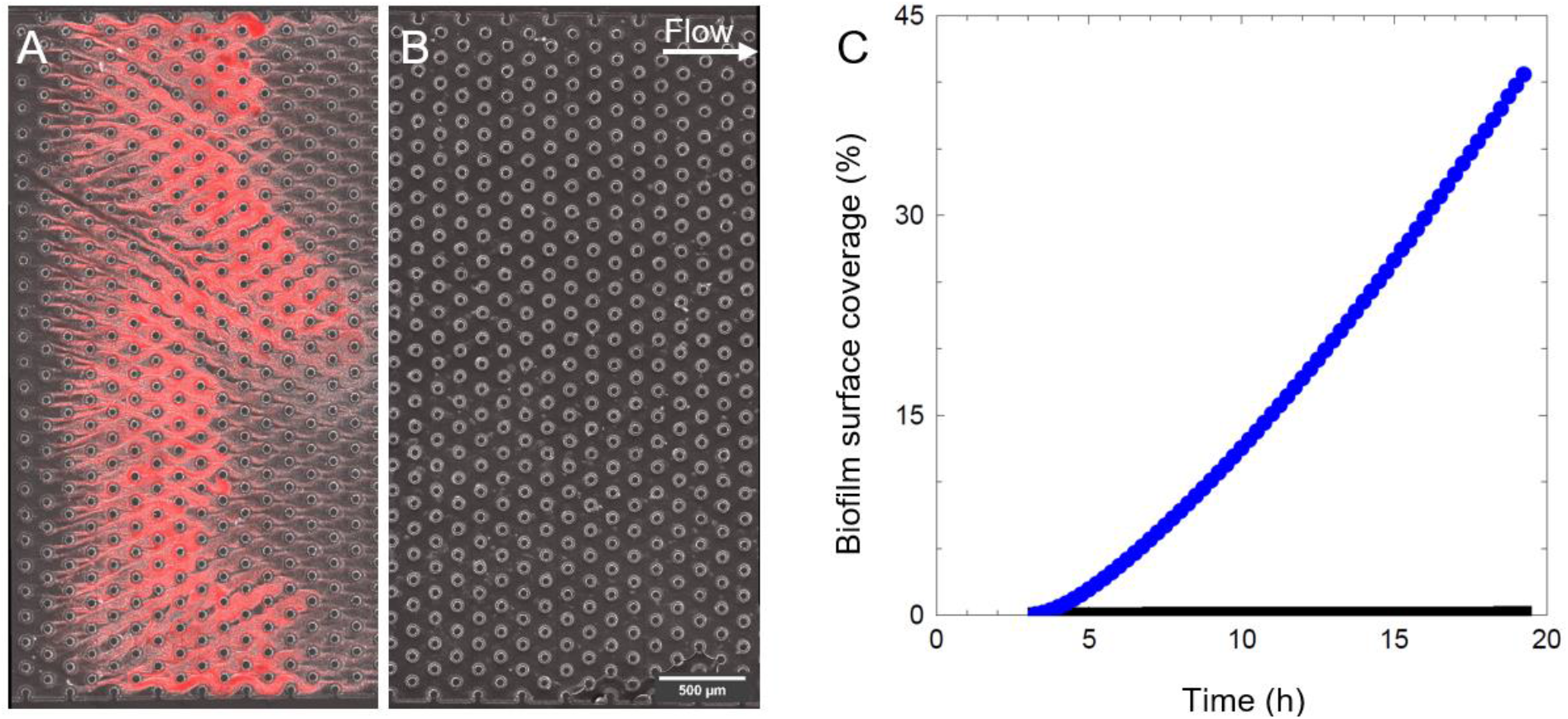
DNase I prevents porous medium clogging by PA14 WT. (*A-B*) Overlaid phase-contrast and fluorescence images of the biofilm formed by *P. aeruginosa* PA14 WT in a model porous medium containing 75-μm pillars, after 23 h of continuous flow of a diluted bacterial suspension (*A*) and of the same suspension containing 1 mg/mL DNase I (*B*), at *U* = 2 mm/s. A portion of the porous medium is shown in the images. (*C*) Biofilm surface coverage, defined as the percentage of the image covered by the biofilm, as a function of time, measured in the same conditions as *A* (blue circles) and *B* (black squares). Biofilm coverage was measured from the fluorescence image and confirmed by comparison with the phase-contrast image.

**Fig. S10.**
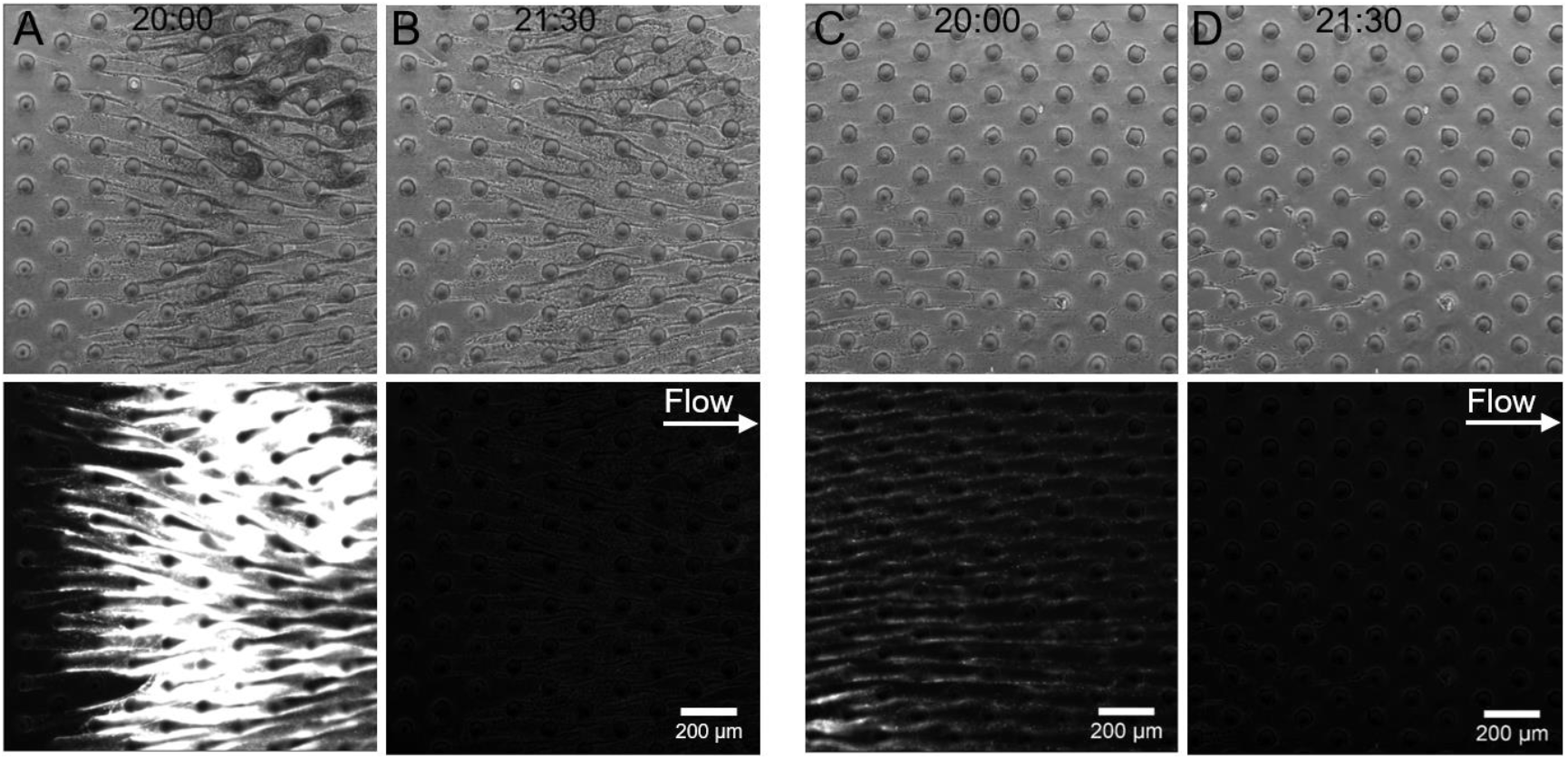
DNase I treatment reduces clogging of porous media by established PA14 WT biofilms. Phase-contrast (top) and red-fluorescence (PI, 2 μg/mL; bottom) images of the biofilm formed by *P. aeruginosa* PA14 WT in a model porous medium after 20 h (*A, C*) of continuous flow of a dilute bacterial suspension at *U =* 2 mm/s, and after 1.5 h flow of 1 mg/mL DNase I in PBS (*B, D*). Two positions in the channel are selected, one in the region clogged by biofilms (*A, B*) and one in the region where just biofilm streamers are present (*C, D*). The treatment depletes the eDNA in the clogged region, as can be seen from the abrupt reduction of the PI signal, and completely clears the region colonized by biofilm streamers.

**Fig. S11.**
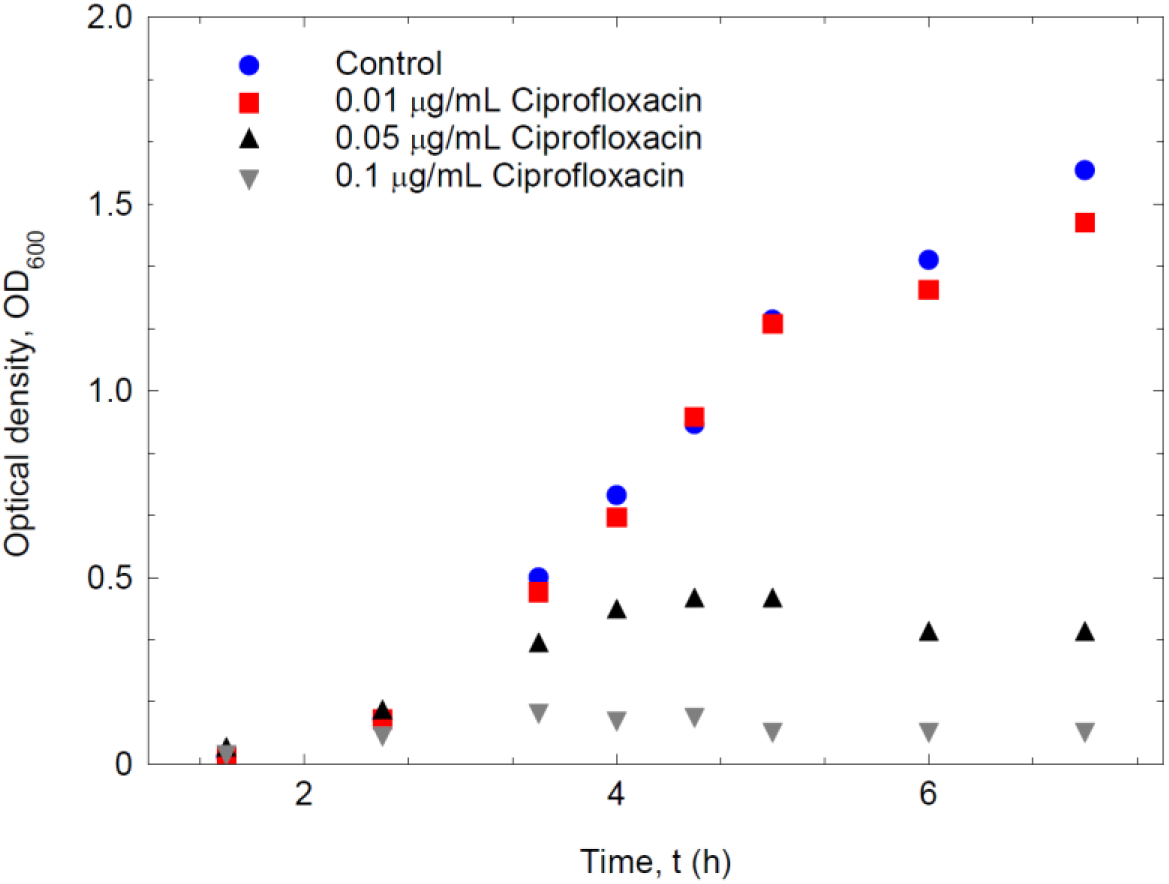
Growth curve for *P. aeruginosa* PA14 WT in the presence or absence of ciprofloxacin. Growth was quantified by measuring the optical density at 600 nm (OD_600_) of *P. aeruginosa* PA14 WT suspensions growing at 37 °C in Tryptone Broth containing different concentrations of ciprofloxacin. Blue circles represent the untreated case. A ciprofloxacin concentration of 0.01μg/ml does not affect bacterial growth (red squares).

**Fig. S12.**
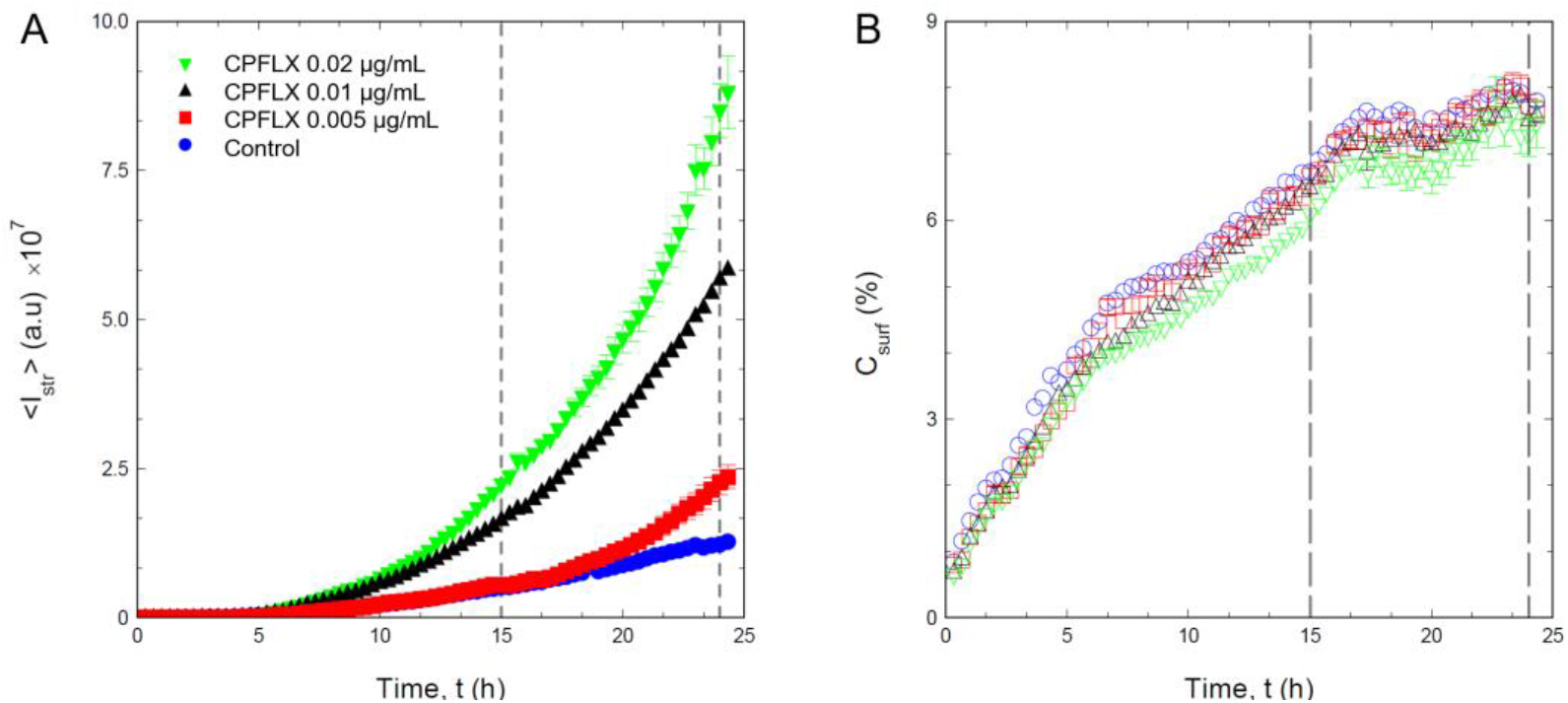
Ciprofloxacin increases streamer growth in PA14 WT. (*A*) Fluorescence intensity of streamers, 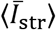, formed by a dilute suspension of *P. aeruginosa* PA14 WT in flow at *U* = 2 mm/s as a function of time, for different concentrations of ciprofloxacin (CPFLX) (blue, no CPFLX; red, 0.005 μg/mL; black, 0.01 μg/mL; green, 0.02 μg/mL). (*B*) Surface coverage (percentage), *C*_surf_, as a function of time, measured on the glass surface 3 mm upstream from the pillar in the phase-contrast images under the same conditions as *A*. Dashed gray lines indicate the time points at which the intensity and surface coverage reported in Fig. 5*E, J* were measured.

**Fig. S13.**
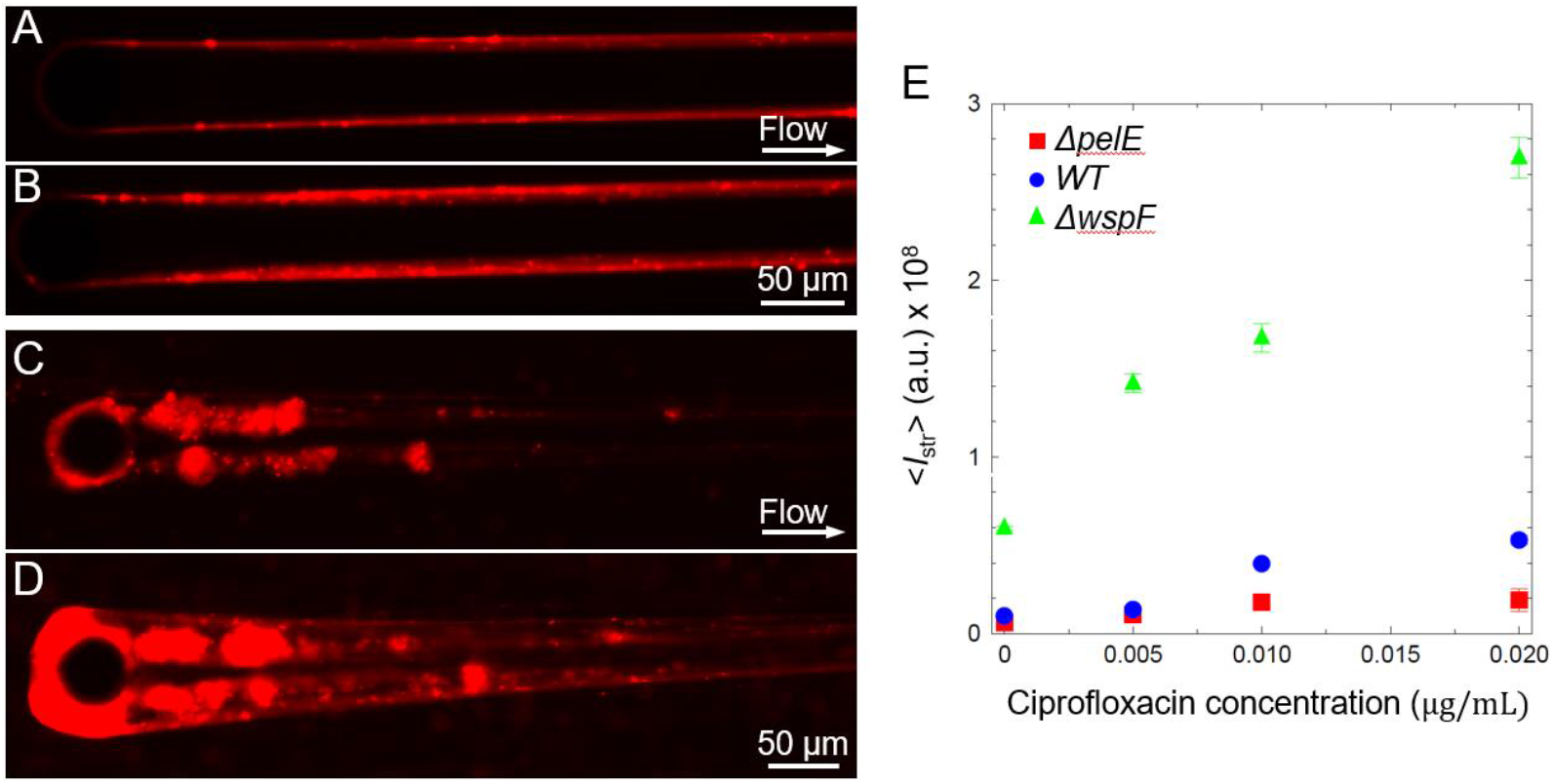
Ciprofloxacin increases streamer growth for both the Pel-deficient strain PA14 *ΔpelE* and Pel-overproducer strain *ΔwspF*. (*A-D*) Fluorescence images of the biofilm streamers formed by *P. aeruginosa* PA14 *ΔpelE* (*A-B*) and *ΔwspF* (*C-D*) attached to a 50-μm pillar after 21 h of continuous flow at *U* = 2 mm/s of a dilute bacterial suspension containing no CPFLX (*A, C*) and ciprofloxacin at a concentration of 0.02 μg/mL (*C, D*). (*E*) Fluorescence intensity of the streamers measured after 21 h of continuous flow at *U* = 2 mm/s, 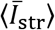, as a function of ciprofloxacin concentration for a suspension of PA14 WT (blue circles), PA14 *ΔpelE* (red squares) and PA14 *ΔwspF* (green triangles). The error bar is calculated as the standard error of the mean.

**Fig. S14.**
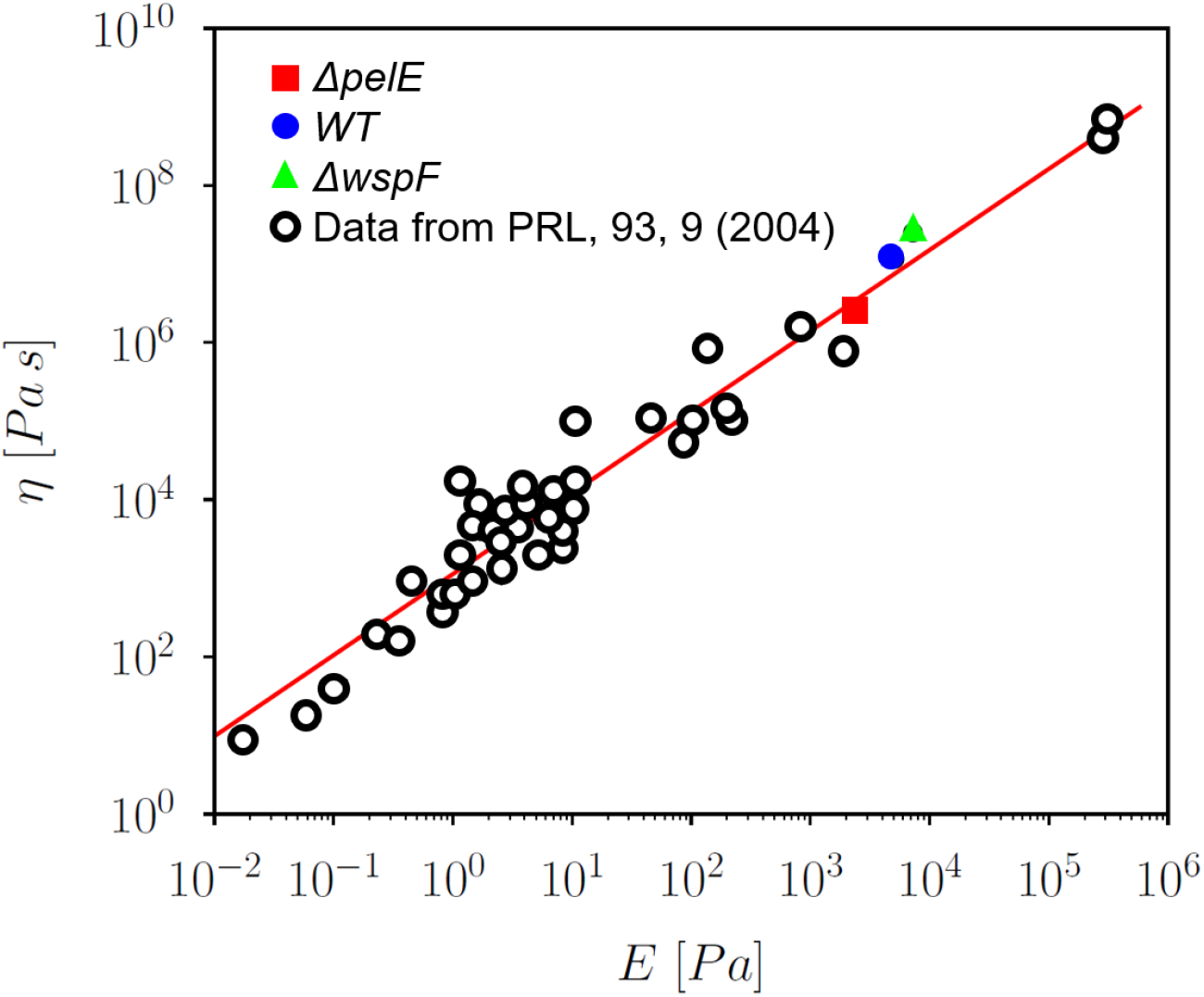
Universal scaling of the elastic relaxation time *η/ E*. Plot of young modulus vs viscosity for data taken from Fig. 3 of (7) (black circles) and for data measured in this work for PA14 WT (blue circles), PA14 *ΔpelE* (red squares) and PA14 *ΔwspF* (green triangles). The straight red line is log *η* = 1.03 log *E* + 1100, also reported in (7). Our data follow the scaling found in (7).

## References

1. H.-C. Flemming, et al., Biofilms: An emergent form of bacterial life. Nat. Rev. Microbiol. 14, 563–575 (2016).

2. H. C. Flemming, S. Wuertz, Bacteria and archaea on Earth and their abundance in biofilms. Nat. Rev. Microbiol. 17, 247–260 (2019).

3. H. Flemming, J. Wingender, The biofilm matrix. Nat. Rev. Microbiol. 8, 623–33 (2010).

4. B. W. Peterson, et al., Viscoelasticity of biofilms and their recalcitrance to mechanical and chemical challenges. FEMS Microbiol. Rev. 39, 234–245 (2015).

5. L. Hall-Stoodley, et al., Towards diagnostic guidelines for biofilm-associated infections. FEMS Immunol. Med. Microbiol. 65, 127–145 (2012).

6. M. Donlan Rodney, J. W. Costerton, Biofilms: Survival mechanisms of clinically relevant microorganisms. Clin. Microbiol. Rev. 15, 167–193 (2002).

7. H. C. Flemming, Biofouling in water systems - Cases, causes and countermeasures. Appl. Microbiol. Biotechnol. 59, 629–640 (2002).

8. P. Desmond, J. P. Best, E. Morgenroth, N. Derlon, Linking composition of extracellular polymeric substances (EPS) to the physical structure and hydraulic resistance of membrane biofilms. Water Res. 132, 211–221 (2018).

9. S. Pandit, et al., The exopolysaccharide component of extracellular matrix is essential for the viscoelastic properties of Bacillus subtilis biofilms. Int. J. Mol. Sci. 21, 6755 (2020).

10. B. Kundukad, et al., Mechanical properties of the superficial biofilm layer determine the architecture of biofilms. Soft Matter 12, 5718–5726 (2016).

11. E. J. Stewart, M. Ganesan, J. G. Younger, M. J. Solomon, Artificial biofilms establish the role of matrix interactions in staphylococcal biofilm assembly and disassembly. Sci. Rep. 5, 13081 (2015).

12. S. G. V. Charlton, et al., Regulating, measuring, and modeling the viscoelasticity of bacterial biofilms. J. Bacteriol. 201, e00101–19 (2019).

13. T. Shaw, M. Winston, C. J. Rupp, I. Klapper, P. Stoodley, Commonality of elastic relaxation times in biofilms. Phys. Rev. Lett. 93, 098102 (2004).

14. E. S. Gloag, S. Fabbri, D. J. Wozniak, P. Stoodley, Biofilm mechanics: Implications in infection and survival. Biofilm 2, 1000017 (2020).

15. L. Hall-Stoodley, J. W. Costerton, P. Stoodley, Bacterial biofilms: From the natural environment to infectious diseases. Nat. Rev. Microbiol. 2, 95–108 (2004).

16. S. C. Chew, et al., Dynamic remodeling of microbial biofilms by functionally distinct exopolysaccharides. MBio 5, e01563–14 (2014).

17. C. D. Nadell, D. Ricaurte, J. Yan, K. Drescher, B. L. Bassler, Flow environment and matrix structure interact to determine spatial competition in *Pseudomonas aeruginosa* biofilms. Elife 6, e21855 (2017).

18. R. Rusconi, S. Lecuyer, N. Autrusson, L. Guglielmini, H. A. Stone, Secondary flow as a mechanism for the formation of biofilm streamers. Biophys. J. 100, 1392–1399 (2011).

19. R. Rusconi, S. Lecuyer, L. Guglielmini, H. A. Stone, Laminar flow around corners triggers the formation of biofilm streamers. J. R. Soc. Interface 7, 1293–1299 (2010).

20. U. U. Ghosh, H. Ali, R. Ghosh, A. Kumar, Bacterial streamers as colloidal systems: Five grand challenges. J. Colloid Interface Sci. 594, 265–278 (2021).

21. D. Scheidweiler, H. Peter, P. Pramateftaki, P. de Anna, T. J. Battin, Unraveling the biophysical underpinnings to the success of multispecies biofilms in porous environments. ISME J. 13, 1700–1710 (2019).

22. K. Drescher, Y. Shen, B. L. Bassler, H. A. Stone, Biofilm streamers cause catastrophic disruption of flow with consequences for environmental and medical systems. Proc. Natl. Acad. Sci. U. S. A. 110, 4345–50 (2013).

23. I. Biswas, M. Sadrzadeh, A. Kumar, Impact of bacterial streamers on biofouling of microfluidic filtration systems. Biomicrofluidics 12, 044116 (2018).

24. N. Debnath, et al., Abiotic streamers in a microfluidic system. Soft Matter 13, 8698–8705 (2017).

25. M. Hassanpourfard, et al., Bacterial floc mediated rapid streamer formation in creeping flows. Sci. Rep. 5, 130070 (2015).

26. S. Das, A. Kumar, Formation and post-formation dynamics of bacterial biofilm streamers as highly viscous liquid jets. Sci. Rep. 4, 7126 (2014).

27. D. J. Wozniak, et al., Alginate is not a significant component of the extracellular polysaccharide matrix of PA14 and PAO1 *Pseudomonas aeruginosa* biofilms. Proc. Natl. Acad. Sci. 100, 7907–7912 (2003).

28. L. Friedman, R. Kolter, Genes involved in matrix formation in *Pseudomonas aeruginosa* PA14 biofilms. Mol. Microbiol. 51, 675–690 (2004).

29. L. Friedman, R. Kolter, Two genetic loci produce distinct carbohydrate-rich structural components of the *Pseudomonas aeruginosa* biofilm matrix. J Bacteriol 186, 4457–4465 (2004).

30. E. E. Mann, D. J. Wozniak, *Pseudomonas* biofilm matrix composition and niche biology. FEMS Microbiol. Rev. 36, 893–916 (2012).

31. M. Fazli, et al., Minireview regulation of biofilm formation in *Pseudomonas* and *Burkholderia* species. Environ. Microbiol. 16, 1961–1981 (2014).

32. B. Konrad, D. Gerd, “Ecology and Epidemiology of Pseudomonas aeruginosa” in Pseudomonas Aeruginosa as an Opportunistic Pathogen, (Springer Science+Business Media, 1993) https:/doi.org/https://doi.org/10.1007/978-1-4615-3036-7_1.

33. J. A. Driscoll, S. L. Brody, M. H. Kollef, The epidemiology, pathogenesis and treatment of *Pseudomonas aeruginosa* infections. Drugs 67, 351–368 (2007).

34. L. K. Jennings, et al., Pel is a cationic exopolysaccharide that cross-links extracellular DNA in the *Pseudomonas aeruginosa* biofilm matrix. Proc. Natl. Acad. Sci. 112, 11353–11358 (2015).

35. M. Klausen, A. Aaes-Jørgensen, S. Molin, T. Tolker-Nielsen, Involvement of bacterial migration in the development of complex multicellular structures in *Pseudomonas aeruginosa* biofilms. Mol. Microbiol. 50, 61–68 (2003).

36. M. J. Franklin, D. E. Nivens, J. T. Weadge, P. Lynne Howell, Biosynthesis of the *Pseudomonas aeruginosa* extracellular polysaccharides, alginate, Pel, and Psl. Front. Microbiol. 2, 1–16 (2011).

37. K. M. Colvin, et al., The Pel and Psl polysaccharides provide *Pseudomonas aeruginosa* structural redundancy within the biofilm matrix. Environ. Microbiol. 14, 1913–1928 (2012).

38. K. M. Colvin, et al., The Pel polysaccharide can serve a structural and protective role in the biofilm matrix of *Pseudomonas aeruginosa*. PLoS Pathog. 7, e1001264 (2011).

39. N. Billings, et al., The extracellular matrix component Psl provides fast-acting antibiotic defense in *Pseudomonas aeruginosa* biofilms. PLoS Pathog. 9(2013).

40. B. J. Cooley, et al., The extracellular polysaccharide Pel makes the attachment of *P. aeruginosa* to surfaces symmetric and short-ranged. Soft Matter 9, 3871–3876 (2013).

41. M. Allesen-holm, et al., A characterization of DNA release in *Pseudomonas aeruginosa* cultures and biofilms. Mol. Microbiol. 59, 1114–1128 (2006).

42. C. B. Whitchurch, T. Tolker-nielsen, P. C. Ragas, J. S. Mattick, Extracellular DNA required for bacterial biofilm formation. Science (80-.). 295, 1487 (2002).

43. T. Seviour, et al., The biofilm matrix scaffold of *Pseudomonas aeruginosa* contains G-quadruplex extracellular DNA structures. npj Biofilms Microbiomes 7 (2021).

44. A. L. Ibáñez de Aldecoa, O. Zafra, J. E. González-Pastor, Mechanisms and regulation of extracellular DNA release and its biological roles in microbial communities. Front. Microbiol. 8, 1390 (2017).

45. L. Montanaro, et al., Extracellular DNA in biofilms. 34, 824–831 (2011).

46. K. C. Rice, et al., The cidA murein hydrolase regulator contributes to DNA release and biofilm development in *Staphylococcus aureus*. Proc. Natl. Acad. Sci. U. S. A. 104, 8113–8118 (2007).

47. U. Böckelmann, et al., Bacterial extracellular DNA forming a defined network-like structure. FEMS Microbiol. Lett. 262, 31–38 (2006).

48. E. A. Izano, M. A. Amarante, W. B. Kher, J. B. Kaplan, Differential roles of poly-N-acetylglucosamine surface polysaccharide and extracellular DNA in *Staphylococcus aureus* and *Staphylococcus epidermidis* biofilms. Appl. Environ. Microbiol. 74, 470–476 (2008).

49. V. Dengler, L. Foulston, A. S. Defrancesco, R. Losick, An electrostatic net model for the role of extracellular DNA in biofilm formation by *Staphylococcus aureus*. J. Bacteriol. 197, 3779–3787 (2015).

50. W. Hu, et al., DNA builds and strengthens the extracellular matrix in Myxococcus xanthus biofilms by interacting with exopolysaccharides. PLoS One 7, e51905 (2012).

51. L. A. Novotny, A. O. Amer, M. E. Brockson, S. D. Goodman, L. O. Bakaletz, Structural stability of Burkholderia cenocepacia biofilms is reliant on eDNA structure and presence of a bacterial nucleic acid binding protein. PLoS One 8, e67629 (2013).

52. Z. Qin, et al., Role of autolysin-mediated DNA release in biofilm formation of *Staphylococcus epidermidis*. Microbiology 153, 2083–2092 (2007).

53. N. Peng, et al., The exopolysaccharide – eDNA interaction modulates 3D architecture of *Bacillus subtilis* biofilm. BMC Microbiol. 20, 115 (2020).

54. L. Tang, A. Schramm, T. R. Neu, N. P. Revsbech, R. L. Meyer, Extracellular DNA in adhesion and biofilm formation of four environmental isolates: A quantitative study. FEMS Microbiol. Ecol. 86, 394–403 (2013).

55. D. M. Dominiak, J. L. Nielsen, P. H. Nielsen, Extracellular DNA is abundant and important formicrocolony strength in mixed microbial biofilms. Environ. Microbiol. 13, 710–721 (2011).

56. Y. Zheng, H. S. Joo, V. Nair, K. Y. Le, M. Otto, Do amyloid structures formed by *Staphylococcus aureus* phenol-soluble modulins have a biological function? Int. J. Med. Microbiol. 308, 675–682 (2018).

57. L. Turnbull, et al., Explosive cell lysis as a mechanism for the biogenesis of bacterial membrane vesicles and biofilms. Nat. Commun. 7, 11220 (2016).

58. J. S. Webb, L. S. Thompson, S. James, T. Charlton, S. Kjelleberg, Cell death in *Pseudomonas aeruginosa* biofilm development. J. Bacteriol. 185, 4585–4592 (2003).

59. W. C. Chiang, et al., Extracellular DNA shields against aminoglycosides in *Pseudomonas aeruginosa* biofilms. Antimicrob. Agents Chemother. 57, 2352–2361 (2013).

60. H. Mulcahy, L. Charron-Mazenod, S. Lewenza, Extracellular DNA chelates cations and induces antibiotic resistance in *Pseudomonas aeruginosa* biofilms. PLoS Pathog. 4 (2008).

61. E. S. Gloag, et al., Self-organization of bacterial biofilm is facilitated by extracellular DNA. Proc. Natl. Acad. Sci. 110, 11541–11546 (2013).

62. L. Eckhart, H. Fischer, K. B. Barken, T. Tolker-Nielsen, E. Tschachler, DNase1L2 suppresses biofilm formation by *Pseudomonas aeruginosa* and *Staphylococcus aureus*. Br. J. Dermatol. 156, 1342–1345 (2007).

63. P. S. Stewart, M. J. Franklin, Physiological heterogeneity in biofilms. Nat. Rev. Microbiol. 6, 199–210 (2008).

64. E. Secchi, et al., The effect of flow on swimming bacteria controls the initial colonization of curved surfaces. Nat. Commun. 11, 2851 (2020).

65. M. D. Brazas, R. E. W. Hancock, Ciprofloxacin induction of a susceptibility determinant in *Pseudomonas aeruginosa*. Antimicrob. Agents Chemoterapy 49, 3222–3227 (2005).

66. C. Haber, D. Wirtz, Shear-induced assembly of λ-phage DNA. Biophys. J. 79, 1530–1536 (2000).

67. M. S. Yeom, J. Lee, The mechanism of the self-assembly of associating DNA molecules under shear flow: Brownian dynamics simulation. J. Chem. Phys. 122, 184905 (2005).

68. A. Devaraj, et al., The extracellular DNA lattice of bacterial biofilms is structurally related to Holliday junction recombination intermediates. Proc. Natl. Acad. Sci. 116, 25068–25077 (2019).

69. J. Sun, et al., Highly stretchable and tough hydrogels. Nature 489, 133–136 (2012).

70. B. H. Schlomann, T. J. Wiles, E. S. Wall, K. Guillemin, R. Parthasarathy, Sublethal antibiotics collapse gut bacterial populations by enhancing aggregation and expulsion. Proc. Natl. Acad. Sci. U. S. A. 116, 21392–21400 (2019).

71. C. Watters, D. Fleming, D. Bishop, K. P. Rumbaugh, Host responses to biofilm. Prog. Mol. Biol. Transl. Sci. 142, 193–239 (2016).

72. H. J. Serrage, M. A. Jepson, N. Rostami, N. S. Jakubovics, A. H. Nobbs, Understanding the matrix: The role of extracellular DNA in oral biofilms. Front. Oral Heal. 2, 1–8 (2021).

73. J. Schindelin, et al., Fiji: An open-source platform for biological-image analysis. Nat. Methods 9, 676–682 (2012).

## SI References

1. C. Haber, D. Wirtz, Shear-induced assembly of λ-phage DNA. Biophys. J. 79, 1530–1536 (2000).

2. M. S. Yeom, J. Lee, The mechanism of the self-assembly of associating DNA molecules under shear flow: Brownian dynamics simulation. J. Chem. Phys. 122, 184905 (2005).

3. C. Hsieh, A. Balducci, P. S. Doyle, An experimental study of DNA rotational relaxation time in nanoslits. 5196–5205 (2007).

4. L. Turnbull, et al., Explosive cell lysis as a mechanism for the biogenesis of bacterial membrane vesicles and biofilms. Nat. Commun. 7, 11220 (2016).

5. M. Allesen-holm, et al., A characterization of DNA release in *Pseudomonas aeruginosa* cultures and biofilms. Mol. Microbiol. 59, 1114–1128 (2006).

6. C. K. Stover, et al., Complete genome sequence of *Pseudomonas aeruginosa* PAO1, an opportunistic pathogen. Nature 406, 959 (2000).

7. T. Shaw, M. Winston, C. J. Rupp, I. Klapper, P. Stoodley, Commonality of elastic relaxation times in biofilms. Phys. Rev. Lett. 93, 098102 (2004).

